# EEG is better when cleaning effectively targets artifacts

**DOI:** 10.1101/2024.06.06.597688

**Authors:** Neil W Bailey, Aron T Hill, Kate Godfrey, M. Prabhavi N. Perera, Nigel C. Rogasch, Bernadette M. Fitzgibbon, Paul B Fitzgerald

## Abstract

Electroencephalography (EEG) is a useful tool to measure neural activity. However, EEG data are usually contaminated with non-neural artifacts, including voltage shifts generated by eye movements and muscle activity, and other artifacts that are less easily characterizable. The confounding influence of artifacts is often addressed by decomposing data into components, subtracting probable artifactual components, then reconstructing data back into the electrode space. This approach is commonly applied using independent component analysis (ICA). Here, we demonstrate the counterintuitive finding that due to imperfect component separation, component subtraction can artificially inflate effect sizes for event-related potentials (ERPs) and connectivity measures, bias source localisation estimates, and remove neural signals. To address this issue, we developed a method that targets cleaning to the artifact periods of eye movement components and artifact frequencies of muscle components. When tested across different EEG systems and cognitive tasks, our results showed that the targeted artifact reduction method is effective in cleaning artifacts while also reducing the artificial inflation of ERP and connectivity effect sizes and minimizing source localisation biases. Our results suggest EEG pre-processing is better when targeted cleaning is applied, as this improves preservation of neural signals and mitigates effect size inflation and source localisation biases that result from approaches which subtract artifact components across the entire time-series. These improvements enhance the reliability and validity of EEG data analysis. Our method is provided in the freely available RELAX pipeline, which includes a graphical user interface for ease of use and is available as an EEGLAB plugin (https://github.com/NeilwBailey/RELAX).

---

Electroencephalography (EEG) provides the ability to non-invasively measure electrophysiological features of neural activity with millisecond (ms) temporal precision. A large field of research has used EEG to examine neural responses to externally presented stimuli by averaging the recorded EEG signal across repeated stimulus presentations, a technique known as the event-related potential (ERP) (Herrmann & Knight, 2001). The ERP has provided a wealth of information about a range of brain functions, including information about which brain regions are likely generating the activity to process specific stimuli when ERP analyses are paired with source localisation techniques (Pires, Leitão, Guerrini, & Simões, 2014). Additionally, research has used measures of connectivity to provide support for suggestions that functional network activity underpins cognitive processes and that impairments in functional connectivity may relate to pathology (Balconi, Gatti, & Vanutelli, 2018; Buzi, Fornari, Perinelli, & Mazza, 2023; Ganzetti & Mantini, 2013; Kundu, Sutterer, Emrich, & Postle, 2013; Miljevic, Bailey, Murphy, Perera, & Fitzgerald, 2023; Talalay, Kurgansky, & Machinskaya, 2018; Tóth et al., 2017). These applications demonstrate the extensive utility of EEG. However, EEG data are almost always contaminated with an array of both biological and non-biological artifacts i.e., voltage shifts in the EEG data that do not originate directly from the brain (Bailey, Biabani, et al., 2023; Croft & Barry, 2000; Muthukumaraswamy, 2013).

Prominent examples of these artifacts are the electrical signals produced by eye movements and blinks (Croft & Barry, 2000). Since the eye contains an electrical current, eye movements produce voltage shifts that are recorded in the EEG signal, and the movement of the eyelid during a blink causes electrical voltages to be conducted from different locations on the eyeball, both of which project voltage shifts to frontal electrodes (Croft & Barry, 2000). Blink artifacts in particular can be much larger in amplitude than the neural activity that researchers aim to measure (Croft & Barry, 2000). Unfortunately, blinks and eye movements can occur in synchronisation with stimuli and the frequency with which a participant blinks can differ between experimental conditions of interest, potentially confounding the results of ERP studies (Huber, Martini, & Sachse, 2023).

While it is less likely for other types of artifacts to synchronize to stimuli, other artifact types may still reduce the signal-to-noise ratio of the ERP estimates. These other artifacts can include: 1) engagement of the frontalis and temporalis muscles situated under frontal and temporal electrodes, which generate voltage shifts that are detected in the EEG as large amplitude, high frequency (>25Hz) activity (Muthukumaraswamy, 2013); 2) low frequency voltage drift caused by changes in impedance at the interface between electrodes and the scalp (sometimes due to perspiration) (Bagheri, Salam, Velazquez, & Genov, 2016; de Cheveigné & Nelken, 2019; Urigüen & Garcia-Zapirain, 2015), and 3) line noise (where electrodes pick up the 50 or 60Hz alternating current from the electrical equipment in the room). Artifacts can also include the contribution of electromagnetic activity generated by the heart, artifacts generated by the participant moving their head, imperfectly connected electrodes (which can cause large and rapid voltage shifts, flat signals, or extremely noisy signals that appear to record no brain activity at all), and other artifacts with less easily identifiable causes (Pion-Tonachini, Kreutz-Delgado, & Makeig, 2019).

To address artifacts in EEG data, many methods have been proposed. Probably the most commonly implemented method over the last 20 years is to use independent component analysis (ICA) to decompose the data into statistically independent components, and then subtract the components that are identified as probable artifacts from the EEG signal. ICA algorithms separate data into components by assuming that data from each electrode represents a weighted sum of the activity from each source, with the weighting dependent on the distance from the source to the recording electrode (Makeig, Bell, Jung, & Sejnowski, 1995). Based on this assumption, ICA algorithms model each independent component by iteratively decomposing the data into the same number of maximally independent components as there are electrodes, through maximization of the statistical independence of each component from each of the other components (Delorme, Palmer, Onton, Oostenveld, & Makeig, 2012; Makeig et al., 1995; Pion-Tonachini et al., 2019). Once data are separated into components (providing a mixing matrix which represents the spatial weights of each component), components that are likely to reflect artifactual sources are typically subtracted from the mixing matrix. This allows ICA to act as a “spatial filter”, where activity from the spatial components that are deemed to reflect artifacts are removed from the data (Delorme, Sejnowski, & Makeig, 2007). After the artifact components are subtracted, the data can then be reconstructed into the original scalp space by multiplying the reduced mixing matrix with the ICA activation data, providing EEG data that are presumed to reflect neural activity free of artifacts (Delorme et al., 2012; Makeig et al., 1995).

Unfortunately, in practice, it is likely that ICA never provides a perfect decomposition of the EEG data into neural and artifact components (Castellanos & Makarov, 2006; Delorme et al., 2007; Fitzgibbon et al., 2016; Islam, Rastegarnia, & Yang, 2016; Pion-Tonachini et al., 2019). There are several reasons for this. First, ICA assumes that the same number of source components are present as there are electrodes measuring the components (Hyvärinen & Oja, 2000). This assumption is unlikely to be true for neural data (Djuwari, Kumar, & Palaniswami, 2006) and is difficult to estimate a priori. Second, the activity from each source is assumed to have a non-Gaussian distribution, an assumption that is also unlikely to be accurate for all neural activity (Matsuda & Yamaguchi, 2022; Metsomaa, Sarvas, & Ilmoniemi, 2014). Finally, ICA assumes that the activity of each source component is statistically independent from all other source components, an assumption that is also unlikely to be true (Chaumon, Bishop, & Busch, 2015; Islam et al., 2016; Metsomaa et al., 2014). In particular, different brain regions are known to be connected, and functional connectivity between brain regions has been shown to be driven by common oscillatory voltage shifts (Chaumon et al., 2015). Additionally, it is likely that multiple brain regions respond to stimuli at the same time, thus providing a temporal dependence between sources (Chaumon et al., 2015). This commonality of response across brain regions has been shown to be associated with a reduction in neural signals when ICA is used to remove transcranial magnetic stimulation-related artifacts from EEG data (Atti, Belardinelli, Ilmoniemi, & Metsomaa, 2024). Given these issues, while ICA has demonstrated utility for removing artifacts from EEG data, it is likely that when ICA is used for artifact cleaning some neural activity is mixed into the artifact components and removed from the data along with the artifact.

Pre-processing methods that remove electrodes and segments of the data showing extreme artifacts prior to providing the data to the ICA algorithm have been suggested to improve ICA’s decomposition (Bailey, Biabani, et al., 2023; Pion-Tonachini et al., 2019). However, it is unclear how much these pre-processing steps help, and a consensus on the optimal approach for rejecting bad electrodes and data segments has not been established (Bailey, Biabani, et al., 2023; de Cheveigne, 2023; Delorme, 2023). In addition to the rejection of bad electrodes and data segments, several other artifact cleaning methods have been suggested to address the unintended reduction of the neural signal and improve artifact cleaning. One early attempted solution was to apply a wavelet transform to every ICA component (an approach termed “wavelet enhanced ICA” [wICA]) (Castellanos & Makarov, 2006). This was intended to remove only the dominant frequencies from each component, which are assumed to reflect artifacts, while simultaneously preserving the neural signal. Our research has suggested the application of wICA to all components is an overly aggressive artifact cleaning strategy that removes considerable amounts of the neural signal from the data as well as the artifacts (Bailey, Biabani, et al., 2023). Later approaches (including our own) have applied the wavelet transform only to components that are deemed likely to reflect artifact components (Bailey, Biabani, et al., 2023; Bailey, Hill, et al., 2023; Issa & Juhasz, 2019; Mammone, La Foresta, & Morabito, 2011). This wavelet enhanced cleaning of only artifact components is intended to remove the artifact contribution (dominant frequencies) from only the artifact components, while preserving neural activity within the artifact component and preserving non-artifact components entirely.

Independent vector analysis (IVA) has also been tested as a potential improvement over the ICA subtraction approach. IVA maximises both the statistical independence of components from other components, and also maximises the statistical dependence of each component within the component (from timepoint to timepoint) (Anderson, Adali, & Li, 2011; Barban, Chiappalone, Bonassi, Mantini, & Semprini, 2021; Chen, Peng, Yu, & Wang, 2017). This has been suggested to improve the component separation, since data at one timepoint within each neural source is assumed to be related to other timepoints within that source (for example, oscillatory activity with repeating voltage shift cycles within a single brain region) (Barban et al., 2021).

Additionally, multi-channel Wiener filtering (MWF) has been proposed to effectively clean EEG data (Somers, Francart, & Bertrand, 2018). To clean EEG data with MWF, the first step involves obtaining templates of EEG data periods that are identified as containing artifacts and periods identified as containing only neural activity. MWF then uses a delay embedded matrix of data from all electrodes in combination with the artifact and clean data templates to obtain a model of the spatio-temporal patterns that characterize the artifacts through a minimization of the mean squared error algorithm (Somers et al., 2018). Since the delay embedding characterizes the temporal as well as spatial aspects of the artifacts, MWF acts as a spatio-temporal filter, in contrast to ICA and IVA which act only as spatial filters. This allows MWF to reduce artifact topographies by taking into account the temporal patterns as well as the spatial patterns of the artifacts (Somers et al., 2018). This has been suggested to effectively clean artifacts while also better preserving neural activity (Somers et al., 2018).

Another component-based artifact reduction method, denoising source separation (DSS), involves using principal component analysis and a bias filter to obtain components that enhance the power from the neural signal of interest and reduce the power from the noise sources (which includes artifacts) (de Cheveigné & Parra, 2014). These components are ranked by the ratio of the signal power to the noise power, and components below a specific signal-to-noise ratio can be deleted. This approach has been suggested to enhance signal-to-noise ratios while relying less on potentially inconsistent modelling (as is the case for the blind source separation methods such as ICA, wICA, and IVA) (de Cheveigné & Parra, 2014).

Finally, regression approaches are used to subtract the beta weights of eye movement artifacts from each electrode (Croft & Barry, 2000). These regression methods have the benefit of not modelling putative source components prior to subtracting artifact components. However, while regression methods avoid the issues with imperfect component separation, it is still possible that the beta weight subtraction removes neural activity from the data (or incompletely reduces the artifact).

Ideally, the efficacy of each of these methods at removing artifacts while preserving neural activity would be tested using a ground-truth dataset. However, a ground-truth is impossible to determine in real EEG recordings, and while simulation approaches may be useful, it is unclear how closely simulated data reflect real data (Muthukumaraswamy, 2013). As such, a common approach to demonstrate good EEG cleaning efficacy in real data is to: 1) show that the cleaning method effectively reduces estimates of the amount of artifact remaining in the data after cleaning, and 2) show that the cleaning method also preserves the neural signal, often by comparing two previously validated experimental conditions and determining which cleaning method provides the largest effect sizes for the presumed experimental effect (Bailey, Biabani, et al., 2023; Bailey, Hill, et al., 2023; Barban et al., 2021; Delorme, 2023). Note that success in both of these points is critical, as it is not useful for a method to be very effective at reducing artifacts if that efficacy comes at the cost of severely reducing the neural signals of interest contained in the EEG data.

In the current work, we show that the selection of an EEG cleaning method based only on its ability to produce a maximal effect size in an experimental comparison is a flawed approach. In particular, we demonstrate the counterintuitive finding that the removal of neural signals due to imperfect component separation when using component subtraction-based cleaning methods can inflate ERP effect sizes as well as reducing them. Our results show that this imperfect component separation issue is ubiquitous across component-based cleaning methods, including wICA, MWF, IVA, and DSS, and is not resolved by a regression artifact reduction method. Furthermore, we demonstrate that component-based methods can also inflate measures of connectivity between electrodes and produce source localisation estimates that are inaccurate. To address the issue posed by imperfect component separation, we introduce a targeted version of the wICA method. This new approach applies wICA specifically and only to periods of the eye movement artifact components that are likely to reflect artifacts. It also cleans muscle components using only a 15Hz low-pass filter to remove only the high frequency activity characteristic of muscle activity while preserving any potential neural activity in the component. We test this new approach by assessing the ability for each cleaning method to reduce artifacts, but also to preserve neural signals in periods of the data that are not affected by artifacts. We perform these tests in two large datasets across different EEG recording systems and different cognitive tasks to demonstrate that our novel method avoids distorting artifact free EEG data, while also providing effective artifact cleaning. We have provided this new approach within the freely available RELAX EEG pre-processing pipeline (https://github.com/NeilwBailey/RELAX) which was previously developed by our team (Bailey, Biabani, et al., 2023; Bailey, Hill, et al., 2023).

## Methods

### A targeted blink and muscle cleaning method

We introduce here a new method that specifically targets just the artifact aspects of artifact components identified by ICA. A schematic explanation of the proposed method can be viewed in Figure 1. Step one involves identifying eye movement artifact components. To achieve this, our algorithm uses both the machine learning algorithm ICLabel (Pion-Tonachini et al., 2019) and icablinkmetrics (Pontifex, Miskovic, & Laszlo, 2017). The icablinkmetrics function was implemented in addition to ICLabel as we have found through informal testing that sometimes ICLabel mis-identifies a blink component as a brain component.

**Figure 1.**
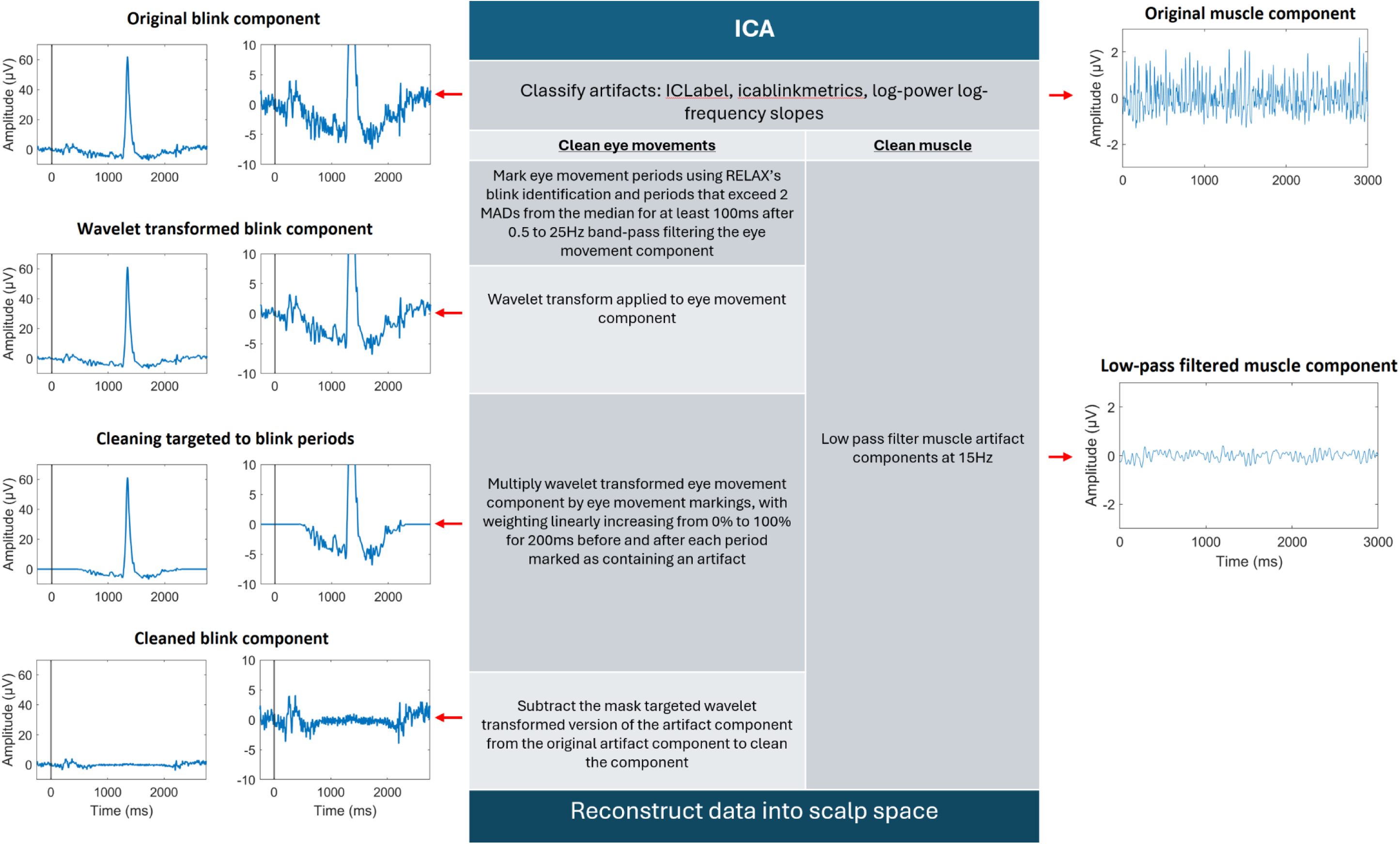
The procedure used to implement the targeted wICA cleaning method, with the effects of this cleaning approach shown for blink related components (left) and muscle related components (right). Within the blink related components, a stimulus presentation is marked with a vertical black line (the images on the far left show a scale that depicts the full blink artifact, while images on the near left show a scale that enables visualisation of the ERP). Note the probable ERP signal immediately after stimulus presentation is preserved by the targeted wICA cleaning approach, whereas a component subtraction approach removes this ERP signal along with the artifacts (wICA – wavelet enhanced independent component analysis, ERP – event related potential).

Step two involves targeting the cleaning of artifact components to be specific to only periods of the component that are affected by eye movement artifacts. To achieve this, our method first bandpass filters each eye movement artifact component from 0.5 to 25Hz using an acausal fourth-order Butterworth filter. This is followed by identification of periods that contain values exceeding 2 median absolute deviations (MAD) from the median for 100ms consecutively, which are annotated as data periods to be corrected. Additionally, the default RELAX blink artifact detection method is used to identify 400ms on either side of a blink maximum as data to be corrected (Bailey, Biabani, et al., 2023; Bailey, Hill, et al., 2023). This creates a temporal mask of the eye movement artifact periods.

Step three involves characterising the dominant frequencies within the eye movement components. To achieve this, a stationary wavelet transform is implemented on the activity from each eye movement component to provide a data driven estimate of the artifact frequencies. This estimate can be subtracted from the original component signal to clean the component (the wICA method performs subtraction of the wavelet transformed artifact model at this point without further modification). To implement the wavelet transform, MATLAB’s “ddencmp” function is used, applying the default settings to automatically set the artifact reduction threshold and automatically determine whether a soft or hard threshold is best suited to characterising the dominant frequencies in the data (Donoho & Johnstone, 1994; Gabard-Durnam, Mendez Leal, Wilkinson, & Levin, 2018). The threshold output from ddencmp is then multiplied by 2 to obtain optimal blink cleaning (a setting that was determined by our iterative testing of the efficacy of different thresholds at both protecting the ERP aspect and cleaning the blink aspect of the components). As per previous implementations of wICA, the wavelet transform is performed on the component using the MATLAB function “swt” with the “coif5” wavelet setting (Bailey, Biabani, et al., 2023; Gabard-Durnam et al., 2018). Then, the stationary wavelet transform with the automated threshold settings are applied to the component using MATLAB’s “wthresh” function (Daubechies, 1992; Gabard-Durnam et al., 2018). The outcome of step three is a model of the full time-series of the eye movement artifact component that represents the dominant frequencies in the signal.

Step four combines the eye movement artifact masks for each eye movement artifact component created in step two with the wavelet transformed versions of each component created in step three. This combination enables the method to target cleaning specifically at eye movement artifact periods, specifically within eye movement artifact components. To achieve this, the wavelet transformed version of each eye movement artifact component is multiplied by a weighted version of the eye movement artifact mask. This weighting progressively increases from 0% during periods that are more than 200ms away from an artifact period to 100% during the periods marked as artifacts, with a linear increase in the weighting from 200ms to 0ms from the artifact marking.

Weighting the wavelet transformed artifact model by the eye movement artifact template in this way ensures no sharp voltage jumps are inserted into the component time-series at the borders of the identified artifact periods, in case the wavelet transformed version of the component and the original component contain values that are offset from each other. Finally, the template weighted wavelet enhanced artifact component (which is targeted to model only artifact periods) is subtracted from the original version of the artifact component. The result of this is a wavelet enhanced ICA cleaning that is targeted to only artifact periods of each eye movement artifact component, which preserves any neural activity mixed into the artifact component outside of artifact periods.

Further, our results indicate that component subtraction cleaning of muscle components also removed ERP data, and that wICA cleaning of muscle components left a considerable amount of muscle activity remaining in the data after cleaning for some files. To address this, our new method uses empirically derived log-frequency log-power slopes reported by Fitzgibbon et al. (2016) to identify muscle components (instead of using ICLabel to identify muscle components). Additionally, instead of subtracting muscle artifact components or using wavelet enhanced cleaning of the muscle artifact components, our novel method simply uses a fourth order acausal Butterworth filter to low-pass filter the muscle components at 15Hz. This reduces the muscle activity contribution to these components, while preserving ERP activity (which is predominantly comprised of activity <15Hz (Schneider & Maguire, 2018)). Finally, since ERP signals were also mixed into other types of artifact components than muscle and eye movements (Supplementary materials Figure S1, Section 3, page 14), and these non-muscle and non-eye movement artifacts are not readily characterizable in a way that would enable targeted reduction, our new method does not remove or reduce these components to avoid distorting the neural signal.

### Comparison of component-based artifact reduction methods

Computations and analyses were performed using Matlab (2023.0, The MathWorks, Inc.) EEGLAB (2023a), fieldtrip (20230328), and JASP (0.17.2.1) (Delorme & Makeig, 2004; Love et al., 2019; Oostenveld, Fries, Maris, & Schoffelen, 2011). To assess the impact of applying component-based artifact reduction methods on ERP data in the context of probable imperfect source component separation, we compared our new method (termed ‘targeted wICA’ hereafter), to eight other source component separation and component reduction methods used to remove artifacts from the data:1) ICA subtract, 2) ICA subtract light, 3) wICA, 4) IVA, 5) MWF, 6) another novel method that we tested: canonical correlation analysis to clean muscle activity followed by generalised eigenvector decomposition to clean blinks (CCA GED, this method is fully explained in the supplementary materials), 7) DSS, and 8) a regression blink reduction method. We note that ICA subtract light differs from ICA subtract in that higher artifact classification thresholds were set in ICLabel before a component was identified as an artifact. In addition to targeted wICA, we developed CCA GED in an initial attempt to address the imperfect source component separation issue by using an alternative component computation approach. This pipeline first applied extended CCA to reduce muscle artifacts (Janani et al., 2018) then applied a GED to reduce blink activity. The GED approach used blink periods to construct a signal covariance matrix and non-blink periods to construct a reference covariance matrix (Cohen, 2022), then performed a GED to decompose data into components reflective of the maximal difference between the blink and non-blink periods. Components that contained blink periods with absolute amplitudes that were significantly larger than non-blink periods (via a one-way t-test of the absolute amplitude of each blink period relative to the absolute amplitude of the overall data) were subtracted from the data before scalp space data were reconstructed. Finally, we tested a traditional regression-based blink cleaning method, which involves subtracting beta weights from each electrode rather than subtracting an artifact component. Full details of these methods are provided in the Supplementary materials (Section 2, page 10).

### Datasets

To demonstrate the optimal pipeline settings across different EEG recording equipment and multiple EEG tasks, we tested our pipelines on the following datasets obtained from healthy participants: 1) A Go/No-go task dataset recorded using a Neuroscan Synamps 2 amplifier (Compumedics, Melbourne, Australia) and 64-channel QuickCap with a sampling rate of 1000Hz and an online 0.01Hz high-pass and 200Hz low-pass filter. The ground electrode was located at AFz and the online reference electrode was located between Cz and CPz. This dataset included 63 participants (23 female and 40 male participants, mean years of age = 33.91, SD = 13.78, range = 20 to 64) (Bailey, Hill, et al., 2023); 2) An N400 word-pair judgement paradigm obtained from the ERPCore dataset (Kappenman, Farrens, Zhang, Stewart, & Luck, 2021). These data were recorded using a 30-channel Biosemi ActiveTwo EEG system with active electrodes, a common mode sense electrode located at PO1 and a driven right leg electrode at PO2. Horizontal electrooculogram (HEOG) electrodes were placed lateral to the external canthus of each eye, and a vertical electrooculogram (VEOG) placed below the right eye. Data were recorded at 1024Hz with a fifth order sinc filter with a half-power cutoff at 204.8Hz in single-ended mode, where voltages were measured between the active and ground electrodes without a reference. The dataset included 40 participants (25 female and 15 male; Mean years of age = 21.50, SD = 2.87, range = 18 to 30). However, we note that only 27 participants provided enough task related epochs that did not contain blinks for both task conditions to enable their inclusion in the analysis of the effect of blink correction on the data from non-blink epochs. Additionally, one further participant was excluded from the blink metric comparisons for only providing two blink related epochs (after extreme outlier exclusions).

Ethics approval for the Go/No-go dataset was provided by the Ethics Committees of Monash University and Alfred Hospital, and ethics approval for the N400 dataset was provided by the Institutional Review Board at the University of California, Davis. All participants provided written informed consent prior to participation in the study.

Prior to testing our novel method and the comparison pipelines, we examined the ERP data without applying any artifact reduction method (but after rejecting extreme outlying periods and channels). This allowed us to determine which ERP periods showed experimental effects prior to any potential adverse effect of a cleaning method, indicating the ERP periods that would be suitable for examination in our analyses. We also tested the effects on ERP comparisons of a range of high-pass filter settings, and a range of extreme artifact rejection approaches and settings. Since these comparisons were made prior to any component-based artifact correction, these tests avoid the risks of component-based artifact reductions on effect size inflation mentioned earlier, so these effect size tests provide a valid approach to determine optimal high-pass filter and extreme artifact rejection settings. A complete explanation of the methods to perform these tests and their results is provided in the Supplementary materials (Section 1). To summarise the results of these tests, a fourth order 0.5Hz high-pass Butterworth filter provided the largest effect sizes for both the P3 and N400 ERPs (compared to 0.25Hz, 0.75Hz and 1Hz settings), so this 0.5Hz high-pass filter setting was used for our primary analyses. Additionally, extreme artifact rejections applied using moderately aggressive RELAX extreme rejection settings showed the best performance for detection of between condition effects for the P3. These moderate settings were also associated with the least distortion of all ERPs by the artifact reduction cleaning steps, so these settings were used in our primary analyses.

### Cleaning Metrics

To evaluate the performance of the different cleaning pipelines, we used two metrics that assessed the severity of artifacts remaining in the data after cleaning. The first artifact focused metric was the frontal blink amplitude ratio (fBAR) (Robbins, Touryan, Mullen, Kothe, & Bigdely-Shamlo, 2020). The blink amplitude ratio is computed by extracting 4000ms epochs surrounding each blink maximum (excluding epochs that contain multiple blinks). Then, all electrodes and epochs are baseline corrected to the average of the first and last 500ms of each epoch. An absolute transform is then performed on each blink epoch, and the mean of the 1000ms period surrounding each blink maximum is obtained. The mean of this blink affected data is then divided by the mean of the absolute transformed data from the first and last 500ms period in the epoch. This value provides the ratio of the absolute amplitude of the blink to the absolute amplitude of surrounding data that is not affected by a blink. Finally, fBAR is obtained by averaging the BAR across the electrodes closest to the eyes (FP1, FPz, FP2, AF3, and AF4 for the Go/No-go dataset, and FP1 and FP2 for the N400 dataset) (Bailey, Biabani, et al., 2023). fBAR values of 1 suggest the blink has been effectively cleaned, with blink periods being no larger in amplitude than non-blink periods. fBAR values below 1 suggest blinks may have been overcleaned, and fBAR values above 1 suggest blinks have not been completely cleaned from the data. However, we note that it is possible that neural activity is larger during a blink period. This possibility has not been formally tested as far as we are aware but, if true, would mean that fBAR values slightly above 1 may reflect optimum cleaning.

Second, we assessed the number of task related epochs that remained in the data after cleaning and showed log-power log-frequency slopes from 7 to 70Hz that exceeded an empirically defined slope threshold of -0.31, which has been shown to be indicative of muscle contamination (Fitzgibbon et al., 2016). This threshold was established by comparing log-power log-frequency slopes extracted from resting EEG recordings under two conditions: 1) where participants either had received a drug that paralysed their scalp muscles, or 2) after they received a control drug (Fitzgibbon et al., 2016). The slope threshold of -0.31 reflected a slope value that was not exceeded by any recording where participants had their scalp paralysed, but was exceeded by recordings that could have been contaminated by muscle activity (where participants did not have their scalp paralysed) (Fitzgibbon et al., 2016). As such, the more epochs with log-power log-frequency slopes above the threshold, the more epochs contaminated by muscle activity are assumed to remain in the data after cleaning (Bailey, Biabani, et al., 2023). For the Go/No-go dataset, we compared the number of epochs showing muscle activity across all pipelines and against a pipeline that used ICA subtract to clean eye movements but did not clean muscle activity (to provide a baseline “no muscle cleaning” comparison). For the N400 dataset, we compared the ICA subtract, targeted wICA, and MWF pipelines against the baseline ICA subtract cleaning of eye movements only comparator. We did not test the number of epochs with muscle activity remaining after cleaning with the regression method, as the regression pipeline is only applicable to cleaning eye movement activity.

### Testing the effects of imperfect source component separation

#### ERP distortion

To assess whether approaches that subtracted eye movement artifact components had an impact on the periods of data that were free from eye movement artifacts, we excluded epochs that contained any identifiable eye movements <u>or</u> high amplitude activity near the ERP windows of interest. We reasoned that if the component-based artifact reduction methods were reducing the artifacts without distorting the non-artifact affected data, then ERPs constructed from cleaned data trials without these artifacts should show identical ERP effects to the raw data. Following the application of each artifact cleaning method, we obtained three difference ERPs, constructed from only epochs that <u>were not</u> contaminated by eye movements: 1) the average ERP difference at a fronto-polar electrode without applying any artifact component subtraction or reduction (referred to as “raw data”, which acts as a pseudo-ground truth); 2) the average ERP difference at a fronto-polar electrode after removal of the eye movement artifact components and reconstruction of the data into the scalp space (providing the effects of cleaning on the ERP difference); 3) the average ERP difference at a fronto-polar electrode produced by reconstructing only the eye movement artifact components back into the scalp space (providing a depiction of the ERP that is contained in the artifact component due to imperfect source component separation). To ensure these ERPs were not contaminated by the blink artifact, we excluded epochs that contained data within 400ms of a blink maximum (as detected by the RELAX pipeline). We additionally excluded epochs from all conditions when the reconstructed “eye movement component only” data contained values at a fronto-polar electrode exceeding 50µV. This amplitude was low enough that we can be confident that difference ERPs were not affected by blink or eye movement artifacts (noting also that both tasks displayed stimuli directly in front of participants, so eye movements were minimal). We then computed the root mean squared error (RMSE) between a fronto-polar electrode ERP constructed from the raw data (data without any component subtraction or reduction applied) and the ERP of the cleaned data from each pipeline within each participant. RMSE values were compared between pipelines using separate ANOVAs for each of the conditions.

We also computed and provide visualisations of the between condition difference ERP following cleaning with each pipeline. This between condition difference ERP provides a visualisation of the potential distortion of experimental effects from comparisons between ERPs across each condition resulting from each artifact component subtraction or reduction method. Given that we excluded all epochs affected by blinks and all higher amplitude epochs that might be likely to contain other eye movement artifacts, an optimal cleaning method would show low RMSE values and no difference between conditions in the data reconstructed from eye movement components only, as well as a difference ERP from the cleaned data that is identical to the raw data difference ERP. Furthermore, an optimal pipeline should also still show effective cleaning of blink periods.

Next, to assess the ability for different pipelines to preserve the ERP topography within the non-blink periods, we compared effect sizes and topographical maps from comparisons between conditions after no cleaning and cleaning by each of the different approaches. To achieve this, we obtained partial eta-squared (ηp²) values for the between condition comparisons using the topographical ANOVA (TANOVA) within RAGU toolbox (Koenig, Kottlow, Stein, & Melie-García, 2011). The TANOVA approach assesses the difference between conditions at all electrodes by computing the global field potential of the difference topography (dGFP) between the two conditions. Conditions that show a consistent activation that consistently differs from the other condition will produce larger topographical differences after averaging across participants, so the dGFP will be larger when the difference across all electrodes is large and consistent across participants (Koenig et al., 2011). Note that this continuous approach avoids the issues with binarization imposed by counting the number of electrodes that exceed a significance value as a measure of the effect size associated with different cleaning approaches (de Cheveigne, 2023). For the Go/No-go data, we tested effects within the P3 window (with data averaged from 315 to 500ms). Due to the sparser electrode distribution in the N400 dataset, we did not use the TANOVA analyses to compare effect sizes from the related/unrelated prime comparisons across the different cleaning pipelines (although we note that to ensure our initial filtering and extreme artifact rejection steps were not biased towards our pipeline or data we had collected, we did perform TANOVA analyses on the N400 dataset to determine the filtering and extreme artifact rejection settings reported in our Supplementary materials).

#### Connectivity

In addition to the ERP analyses, we examined a measure of connectivity between electrodes, as it has been suggested that component subtraction methods can inflate measures of connectivity (Castellanos & Makarov, 2006). To achieve this, we extracted six second data epochs, as epochs of this length have been suggested to be optimal for avoiding edge effects within connectivity estimation (Miljevic, Bailey, Vila-Rodriguez, Herring, & Fitzgerald, 2022). We then computed the debiased weighted phase lag index (dwPLI) from these six second epochs, as this method is robust to the effects of zero phase lag connectivity which is unlikely to reflect physiologically meaningful connectivity (Vinck, Oostenveld, Van Wingerden, Battaglia, & Pennartz, 2011). We focused our analyses on data from the one second period after stimulus presentation and on connectivity between the FPz and Pz electrodes within the 1 to 13Hz range (reflecting the most prominent oscillations in our data). These dwPLI connectivity values were computed and compared within the Go/No-go dataset and a restricted exemplifying set of cleaning pipelines (to save computation time) that included targeted wICA, ICA subtract, MWF, and the raw (eye movement excluded) data for comparison. This analysis tested whether the component-based artifact reduction approaches might also distort connectivity estimates through the projection of the fronto-central maximal activity to the fronto-polar electrodes, as has been indicated by previous research (Castellanos & Makarov, 2006).

#### Source localization

Finally, within the same restricted set of pipelines, we examined the source localisation estimated dipole locations for the single most prominent dipole at the peak of the averaged Go P3 and No-go P3 ERPs (400ms) within each participant. This was undertaken to test whether the component subtraction approach might also distort estimates of the generators of brain activity. We note that the activity found in late-latency ERPs like the P3 are thought to be generated by multiple dipoles, but here we focus here on the single best fitting dipole as a representative example to enable meaningful analysis of whether different artifact reduction methods distort the results of source localisation. We did not implement this analysis for the N400 dataset due to the small number of electrodes, which limits source localisation accuracy. We achieved source localisation using the EEGLAB function dipfit_erpeeg with the standard boundary element model based on the Montreal Neurological Institute (MNI) canonical brain and exact low-resolution electromagnetic tomorgraphy (eLORETA) to construct the leadfield, enabling estimation of a single best fitting dipole (based on the least residual variance) to each participant’s P3 ERP (Oostendorp & Van Oosterom, 1989; Oostenveld et al., 2011). The MNI Y axis location value for this single best fitting dipole (representing the anterior/posterior location) was extracted to determine whether removing eye movement components led to a dipole source estimate that was more posterior than was the case for the data without component-based artifact reduction.

With the exception of post-hoc tests, which were controlled via Bonferonni-Holm multiple comparison controls (indicated as pHolm values), no multiple comparison controls were applied across our analyses, as we emphasized sensitivity to detect meaningful differences between the pipelines over the risk of false positive results from multiple comparisons (Bailey et al., 2023).

## Results

### Component subtraction distorts artifact-free ERPs

To demonstrate the problem posed by artifact component subtraction methods, we have provided visualizations of different aspects of a typical ICA subtract cleaning approach in Figures 2 to 4. To provide these depictions, the Go/No-go data were cleaned by decomposing data with the Preconditioned ICA for Real Data (PICARD) algorithm (Frank, Makeig, & Delorme, 2022), then applying ICLabel to identify only eye-movement artifact components, which were subtracted from the data before data were reconstructed back into the scalp space. We provide first a depiction of the scalp space topography reconstructed from only the eye-movement components from each participant, averaged for 800ms around each blink across all epochs and all participants. This demonstrates the expected blink artifact topography, with positive voltages in fronto-polar electrodes (Figure 2A). Next, topographies from the same reconstructed eye-movement component artifacts only data are provided (Figure 2B and 2C). However, these topographies were constructed from Go and No-go stimuli locked periods obtained by excluding all epochs containing probable eye movement artifacts (as described in our methods), followed by averaging across all epochs and all participants within the P3 window (315 to 500ms). These topographies enable visualisation of the topographical activity pattern that was removed from the Go and No-go P3 by the eye-movement component subtraction. These topographies show a similar but inverted topographical pattern to the blink-locked topographies, with negative voltages in fronto-polar electrodes. This demonstrates how subtracting eye-movement components from the data can result in alterations to activity in fronto-polar electrodes (and to some extent frontal electrodes) even in task-relevant periods that are not affected by eye-movements, suggesting over-cleaning of the data.

**Figure 2.**
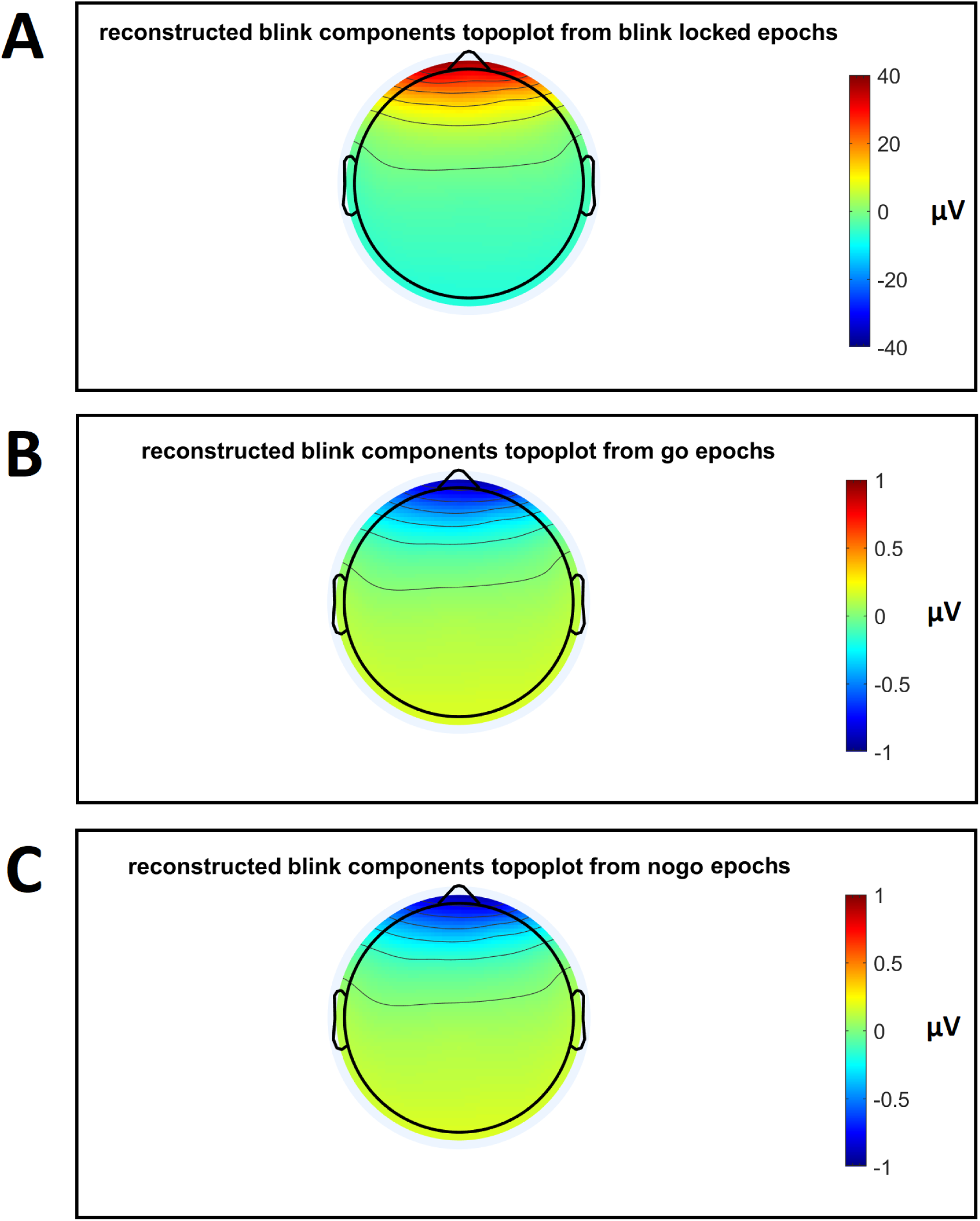
Topographies after reconstructing scalp space data from ICA components after removal of all components except the eye-movement components from each participant. A: The topography averaged for 800ms around each blink across all epochs and all participants. This demonstrates the expected blink artifact topography, with positive voltages in fronto-polar electrodes. B and C: The topography constructed after the exclusion of epochs containing blink activity and epochs where the reconstruction from the “eye movement component only” data contained values at a fronto-polar electrode exceeding 50µV, then averaged across all epochs and all participants within the P3 window (315 to 500ms) following Go (B) and No-go trials (C). Note the negative voltage values at frontal electrodes, demonstrating that ERP topographies are altered by eye-movement component subtraction even when the ERPs are obtained from epochs after eye-movement affected epochs are excluded.

Next, Figures 3A-3G show the effect of this over-cleaning on the Go and No-go ERP trace at FPz and Fz. The ERP pattern that was inadvertently mixed into the eye movement components (and subtracted from the data by the ICA subtract method) is depicted in green and shows a negative voltage deflection during the P3 window. This negative voltage deflection during the P3 window is particularly prominent in the No-go trials. The effect of this component subtraction on the Go and No-go ERPs at FPz is depicted (Figures 3A and 3B), with the raw data (in red) showing a more negative voltage deflection during the P3 window than the cleaned data (in blue), due to the removal of the negative values contained within the eye-movement components (since subtraction of a negative value is equivalent to addition of a positive value). In contrast, the eye-movement component data reconstructed at Fz contained very little ERP activity mixed into the artifact component, and so ERPs at Fz were only minimally distorted (Figures 3C and 3D). The effect of component subtraction cleaning on average amplitudes within the P3 window compared to the raw data are depicted in Figures 3E and 3F, where the difference between the Go and No-go P3 amplitude is larger after ICA subtract was applied to remove eye-movement components. At this point, it is worth highlighting that these data were all extracted from epochs that did not contain eye-movement artifacts, so no difference should be present between the cleaned and raw data.

**Figure 3.**
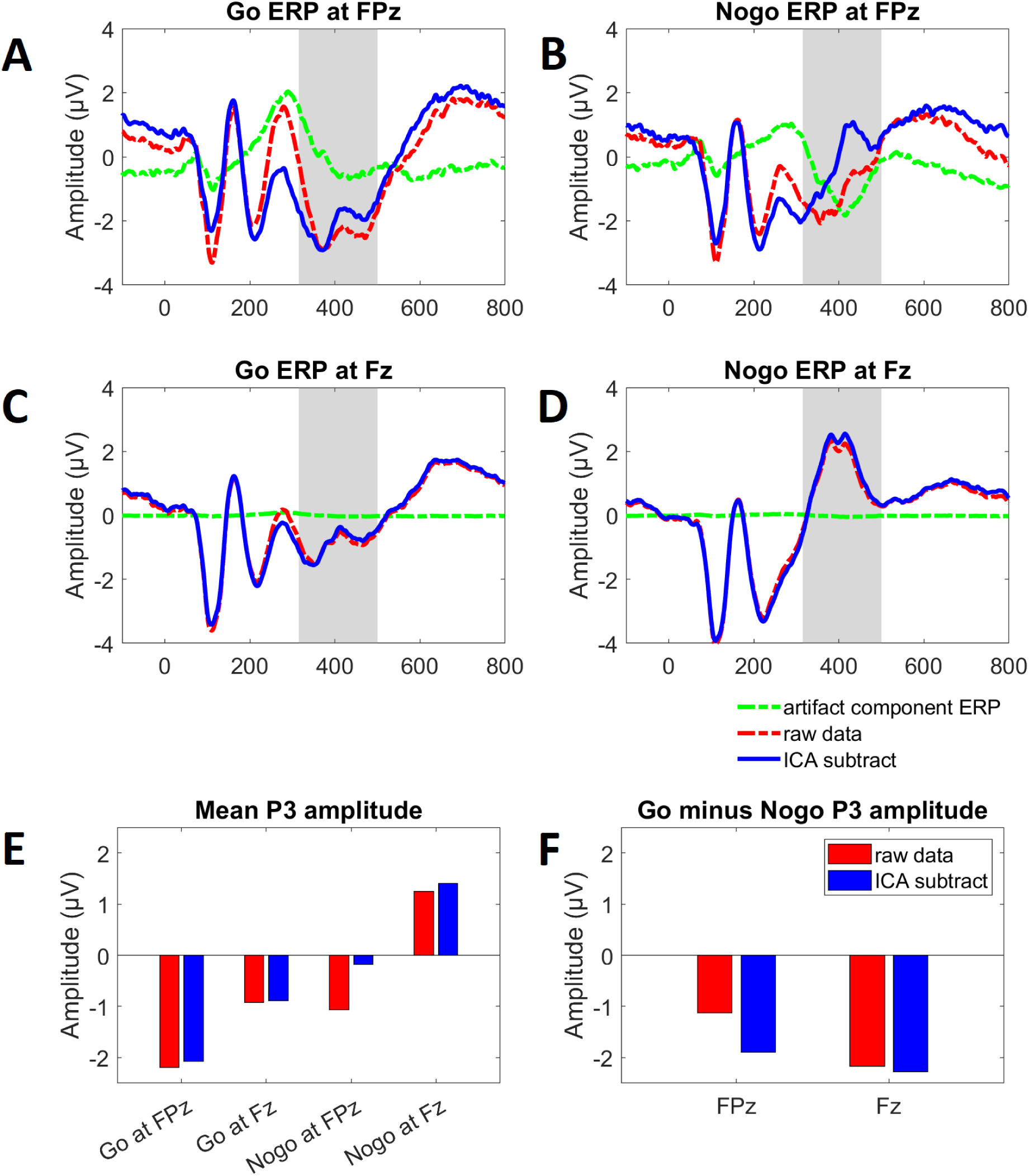
Event-related potential waveforms at FPz (A and B) and Fz (C and D), representing eye-movement components reconstructed into the component space (green), the raw data (before cleaning with ICA subtract - red), and the data after cleaning with ICA subtract (blue). Note that these ERPs were constructed after the exclusion of epochs containing any probable eye movement artifacts, then averaging across all epochs and all participants. Grey shading is provided to indicate the P3 window (315 to 500ms), with plot E showing the averaged amplitude within the P3 window following Go and No-go trials at FPz and Fz and plot F showing the difference between mean Go and No-go P3 amplitudes at these two electrodes. Note the smaller mean amplitude of the No-go P3 at FPz after cleaning with ICA subtract (plot E), which resulted in a larger Go minus No-go difference at FPz (plot F). Paired samples t-tests indicated that this adverse effect of cleaning led to a larger effect size for a comparison between the Go and No-go P3 at FPz in the cleaned data: t(63) = 8.298, pHolm < 0.001, Cohen’s d = 1.051, compared to the raw data: t(63) = 4.952, p < 0.001, Cohen’s d = 0.627. Furthermore, while the Go P3 did not differ between the cleaned and raw data t(63) = 0.877, p < 0.001, Cohen’s d = 0.069, the No-go P3 showed a significant difference between the cleaned and raw data t(63) = 6.272, p < 0.001, Cohen’s d = 0.492. These results show that ERPs measured in a fronto-polar electrode are altered by eye-movement component subtraction even when the ERPs are obtained from epochs after eye-movement affected epochs are excluded.

**Figure 4.**
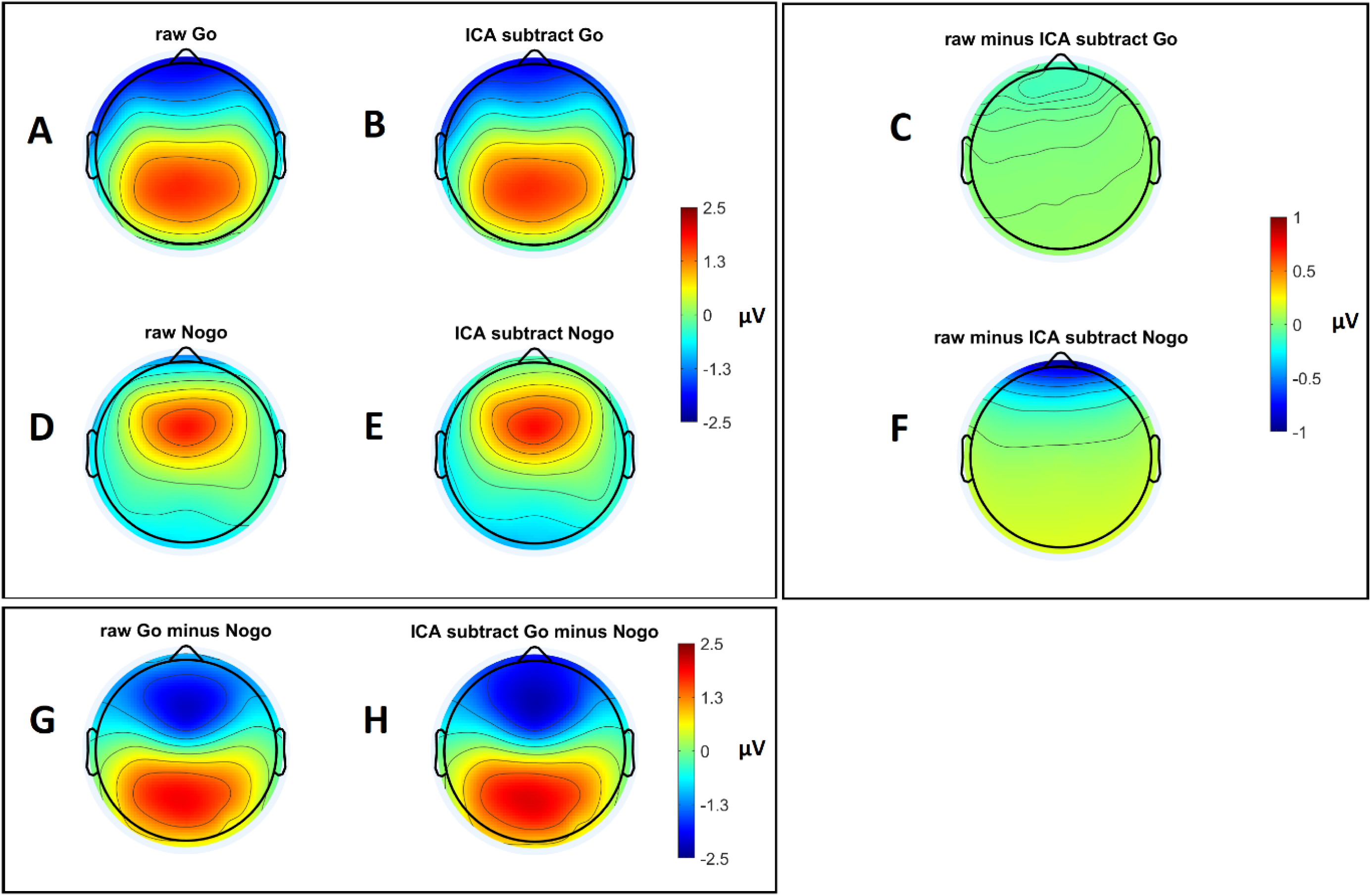
Topographical plots depicting activity averaged within the P3 window (315 to 500ms) across all epochs and participants (A, B, D, and E), obtained after the exclusion of all epochs containing eye movement artifacts. Subplots G and H depict the Go minus No-go P3 topographies. A subtle increase in the strength of negative values for this Go minus No-go difference topography can be seen at fronto-polar electrodes after eye-movement artifacts were cleaned using ICA subtract compared to the raw data. Plots C and F depict the difference between the raw and cleaned data for the Go and No-go P3 respectively. These plots highlight that the difference between the raw and cleaned data present at fronto-polar electrodes is much more prominent in the No-go trials. Due to this difference, the effect size for the TANOVA analysis including all electrodes of the Go vs No-go P3 provided by the cleaned data was larger than the effect size provided by the raw data (p < 0.001, ηp² = 0.390 and p < 0.001, ηp² = 0.332 respectively).

Finally, Figure 4 depicts the Go and No-go topographies, as well as Go minus No-go difference topographies for the P3 window from both the raw and the cleaned data. While the topographies appear to be mostly similar, careful inspection of the difference topographies (Figures 4G and 4H) shows more negative fronto-polar voltages in the ICA subtract Go minus No-go topography than the raw Go minus No-go topography. This is highlighted in the topography shown by Figure 4F, which depicts the difference between the raw and ICA subtract No-go P3 topographies, where the effect of cleaning is most apparent in No-go trials at the fronto-polar electrodes. In contrast, the difference between the raw and ICA subtract Go P3 topographies shows very little difference (Figure 4C). Critically, an ANOVA testing the interaction between the clean and raw data and the Go and No-go trials demonstrated a significant effect (F(1,61) = 25.304, p < 0.001, ηp² = 0.287, ηG² = 0.011). Post-hoc paired samples t-tests indicated that the effect of cleaning led to a larger effect size for the comparison between the Go and No-go P3 at FPz in the cleaned data: t(63) = 8.298, pHolm < 0.001, Cohen’s d = 1.051, compared to the raw data: t(63) = 4.952, p < 0.001, Cohen’s d = 0.627. Furthermore, while the Go P3 did not differ between the cleaned and raw data t(63) = 0.877, p < 0.001, Cohen’s d = 0.069, the No-go P3 showed a significant difference between the cleaned and raw data t(63) = 6.272, p < 0.001, Cohen’s d = 0.492. The difference in effect sizes between the raw and cleaned was also present in the TANOVA analyses that included all electrodes, where the effect size provided by the cleaned data was larger than the effect size provided by the raw data (p < 0.001, ηp² = 0.390 and p < 0.001, ηp² = 0.332 respectively). Together, these results demonstrate that removing independent components representing eye blinks can distort EEG signals in data periods free from eye movements, leading to inflated effect sizes when comparing ERPs between conditions, both at the single electrode level and in comparisons that include all electrodes. In the following sections, we compare a range of different artifact reduction methods, including our novel targeted wICA approach, to identify a cleaning pipeline which best minimises this issue.

### Artifact Cleaning Metrics

With regards to the cleaning of artifacts, our results indicated that all pipelines provided reasonable blink artifact reduction (Table 1 and Figure 5), indicated by mean fBAR values of between 1.04 and 1.40 (in contrast to fBAR values of >3 which are common in raw data). However, the ANOVA indicated that within the Go/No-go dataset there was a significant difference in fBAR values between the different cleaning pipelines (F(7, 63) = 66.728, p < 0.001, ηp² = 0.514, ηG² = 0.278). Post-hoc tests indicated that the targeted wICA and MWF pipelines showed better blink cleaning performance than all other pipelines (all pHolm < 0.001) but did not differ from each other (pHolm = 1.0). DSS showed significantly worse performance than all other pipelines (all pHolm < 0.001), while the remaining pipelines did not differ from each other (all pHolm > 0.9). One caveat worth noting is that the amplitude of blink artifact reduction achieved by targeted wICA within blink periods is identical to the default wICA, so the improved performance of targeted wICA over the default wICA approach must be due to improved preservation of the signal amplitudes outside of blink periods (which are used as a baseline in the fBAR calculation). This preservation of neural signals meant that absolute amplitudes outside of the blink periods were almost identical to the absolute amplitude within blink periods after cleaning with targeted wICA. It also meant that the absolute amplitudes outside of the blink periods after cleaning with targeted wICA were larger than the absolute amplitudes outside of the blink periods after cleaning with the other pipelines due to the improved preservation of non-eye-movement artifact signals.

**Figure 5.**
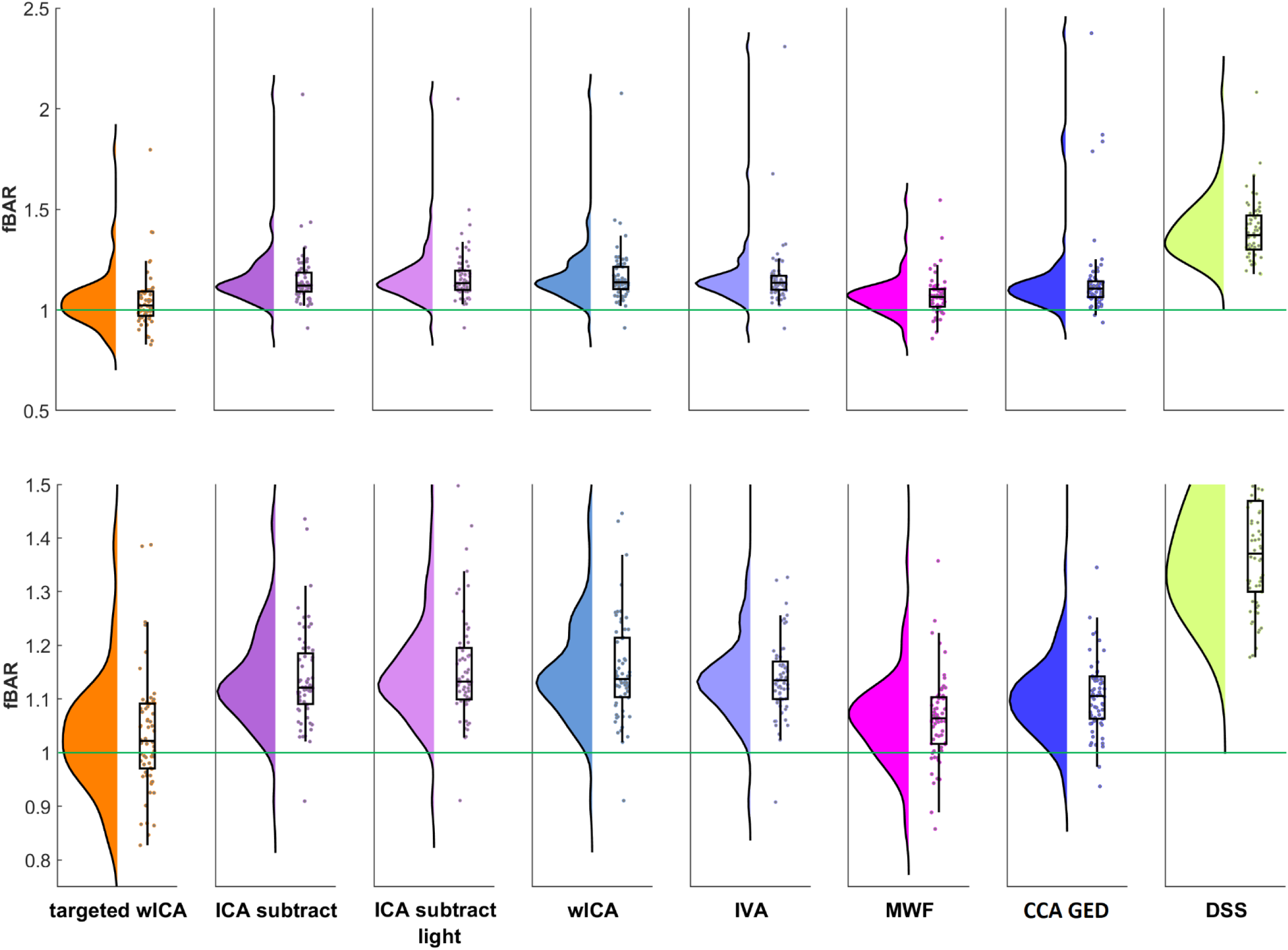
Frontal blink amplitude ratio (fBAR) values for each of the cleaning pipelines demonstrating the efficacy of blink artifact cleaning from each pipeline when applied to the Go/No-go dataset. Above: fBAR values pipeline on a scale that shows all datapoints. Below: fBAR values on a reduced scale so differences between pipelines can be more easily visualised. The green line reflects an fBAR of 1, which would indicate that the mean absolute amplitude within the period affected by the blink is the same as the mean absolute amplitude outside of the period affected by the blink. ICA – independent component analysis, wICA – wavelet enhanced ICA, IVA – independent vector analysis, MWF – multi-channel Wiener filtering, CCA GED – canonical correlation analysis and generalised eigenvector decomposition, DSS – denoising source separation.

**Table 1.** Mean frontal blink amplitude ratio (fBAR) after cleaning with each of the pipelines for the Go/No-go dataset. ICA – independent component analysis, wICA – wavelet enhanced ICA, IVA – independent vector analysis, MWF – multi-channel Wiener filtering, CCA GED – canonical correlation analysis and generalised eigenvector decomposition, DSS – denoising source separation.

| Pipeline | Mean fBAR (SD) |
| --- | --- |
| targeted wICA | 1.044 (0.146) |
| ICA subtract | 1.152 (0.146) |
| ICA subtract light | 1.169 (0.148) |
| wICA | 1.167 (0.148) |
| IVA | 1.164 (0.175) |
| MWF | 1.071 (0.100) |
| CCA GED | 1.156 (0.229) |
| DSS | 1.394 (0.149) |

Similarly, our results indicated that most pipelines provided a somewhat effective muscle artifact reduction, indicated by a mean number of muscle contaminated stimulus locked epochs after cleaning of between 0 and 30 compared to a mean of 40 muscle contaminated epochs when no muscle artifact reduction method was applied (Table 2). The ANOVA indicated there was a significant difference between the cleaning pipelines in muscle artifact reduction performance: (F(7, 63) = 19.482, p < 0.001, ηp² = 0.236, ηG² = 0.167). Post-hoc tests indicated that wICA showed the worst muscle artifact cleaning, worse than all other pipelines (pHolm < 0.001) and not significantly different from the data without cleaning (pHolm = 0.216). This was followed by MWF, which showed a trend towards worse performance than IVA (pHolm = 0.078). All other pipelines did not differ from each other (all pHolm > 0.23). However, we also note that all pipelines except IVA and targeted wICA provided sub-optimal muscle artifact cleaning for at least one of the files, with the least effectively cleaned file by each of these other pipelines showing ≥30 muscle affected epochs. This was particularly bad for the DSS, MWF, ICA, and wICA pipelines. In particular, the default version of wICA seems to have increased number of epochs with log-frequency log-power slopes indicative of muscle activity beyond that found in the raw data for at least one participant (with 195 epochs showing log-frequency log-power values indicative of muscle activity remaining after cleaning). In contrast, the maximum number of epochs showing log-frequency log-power slopes indicative of muscle activity remaining after cleaning was 16 for targeted wICA, and 1 for IVA.

**Table 2.** Mean number of epochs containing log-frequency log-power slopes indicative of muscle activity remaining in the Go/No-go dataset after cleaning with each of the pipelines. ICA – independent component analysis, wICA – wavelet enhanced ICA, IVA – independent vector analysis, MWF – multi-channel Wiener filtering, CCA GED – canonical correlation analysis and generalised eigenvector decomposition, DSS – denoising source separation.

| Pipeline | Mean number of epochs<br>showing muscle artifact after<br>cleaning (SD) | Maximum |
| --- | --- | --- |
| ICA subtract only cleaning eye<br>movements | 40.219 (49.459) | 177 |
| targeted wICA | 1.438 (2.938) | 16 |
| ICA subtract | 4.031 (13.697) | 99 |
| ICA subtract light | 12.172 (26.288) | 116 |
| wICA | 29.078 (54.168) | 195 |
| IVA | 0.016 (0.125) | 1 |
| MWF | 12.859 (32.121) | 136 |
| CCA GED | 2.781 (5.119) | 30 |
| DSS | 9.313 (16.279) | 83 |

Similar results were apparent for the N400 dataset, where our analysis of a restricted number of pipelines indicated that all the tested pipelines provided reasonable blink artifact reduction, indicated by mean fBAR values of between 0.944 and 1.254 (Table 3 and Figure 6). Overall, the ANOVA indicated there was a significant difference between the pipelines (F(3, 26) = 41.357, p < 0.001, ηp² = 0.521, ηG² = 0.339). Post-hoc tests indicated that targeted ICA showed better performance than the other pipelines included in this analysis (ICA subtract, MWF, and regression, all pHolm < 0.001). The MWF and ICA subtract methods did not differ from one another (pHolm = 0.599), and the regression method performed worse than all other pipelines (all pHolm < 0.001).

**Figure 6.**
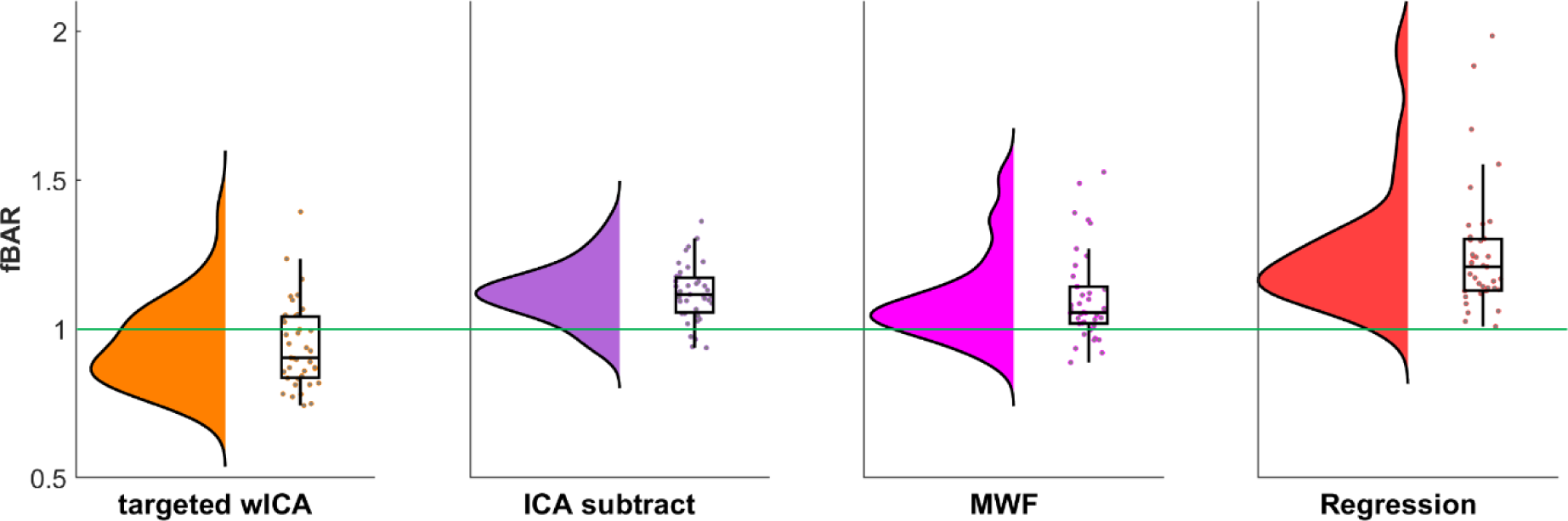
Frontal blink amplitude ratio (fBAR) values for each of the cleaning pipelines demonstrating the efficacy of blink cleaning from each when applied to the N400 dataset. The green line reflects an fBAR of 1, which would indicate that the mean absolute amplitude within the period affected by the blink is the same as the mean absolute amplitude outside of the period affected by the blink. ICA – independent component analysis, wICA – wavelet enhanced ICA, MWF – multi-channel Wiener filtering.

**Table 3.**
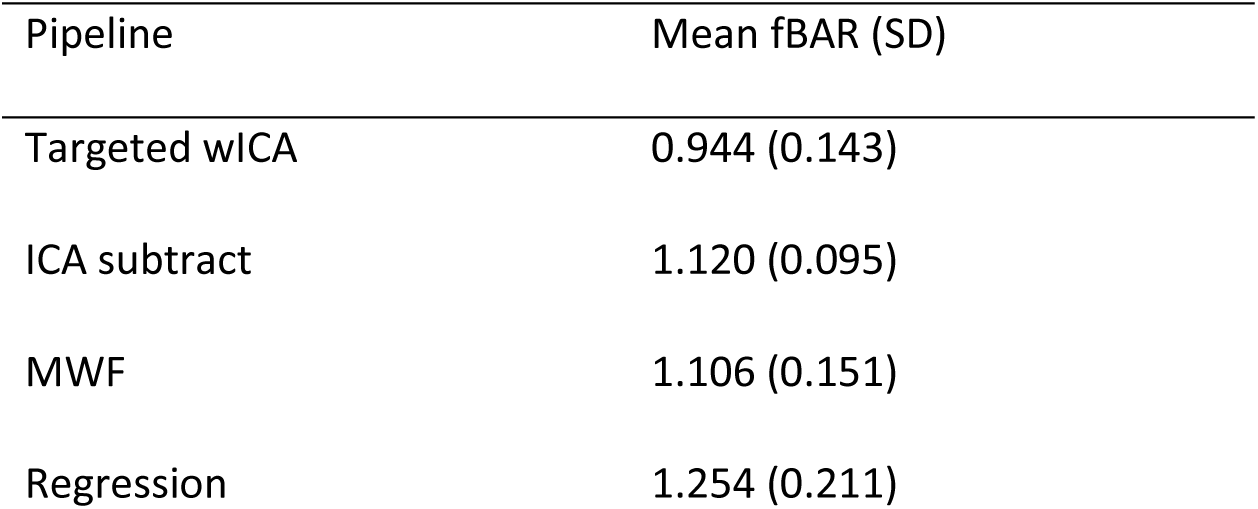
Mean frontal blink amplitude ratio (fBAR) within the N400 dataset after cleaning with each of the pipelines. ICA – independent component analysis, wICA – wavelet enhanced ICA, MWF – multi-channel Wiener filtering.

With regards to the cleaning efficacy for the muscle artifact in the N400 dataset, the ANOVA indicated there was a significant difference between pipelines (F(3, 26) = 4.931, p = 0.003, ηp² = 0.107, ηG² = 0.027, Table 4). Post-hoc tests indicated that ICA subtract and targeted wICA showed the best muscle artifact cleaning, both showing better performance than the baseline comparator pipeline that used ICA subtract to clean only eye movement artifacts (which did not clean muscle activity) (pHolm = 0.004 and 0.015 respectively), but not differing from each other or MWF (all pHolm > 0.245). In contrast, MWF did not show a difference compared to the baseline comparator pipeline that used ICA subtract to clean eye movements but not clean muscle activity (pHolm = 0.350).

**Table 4.**
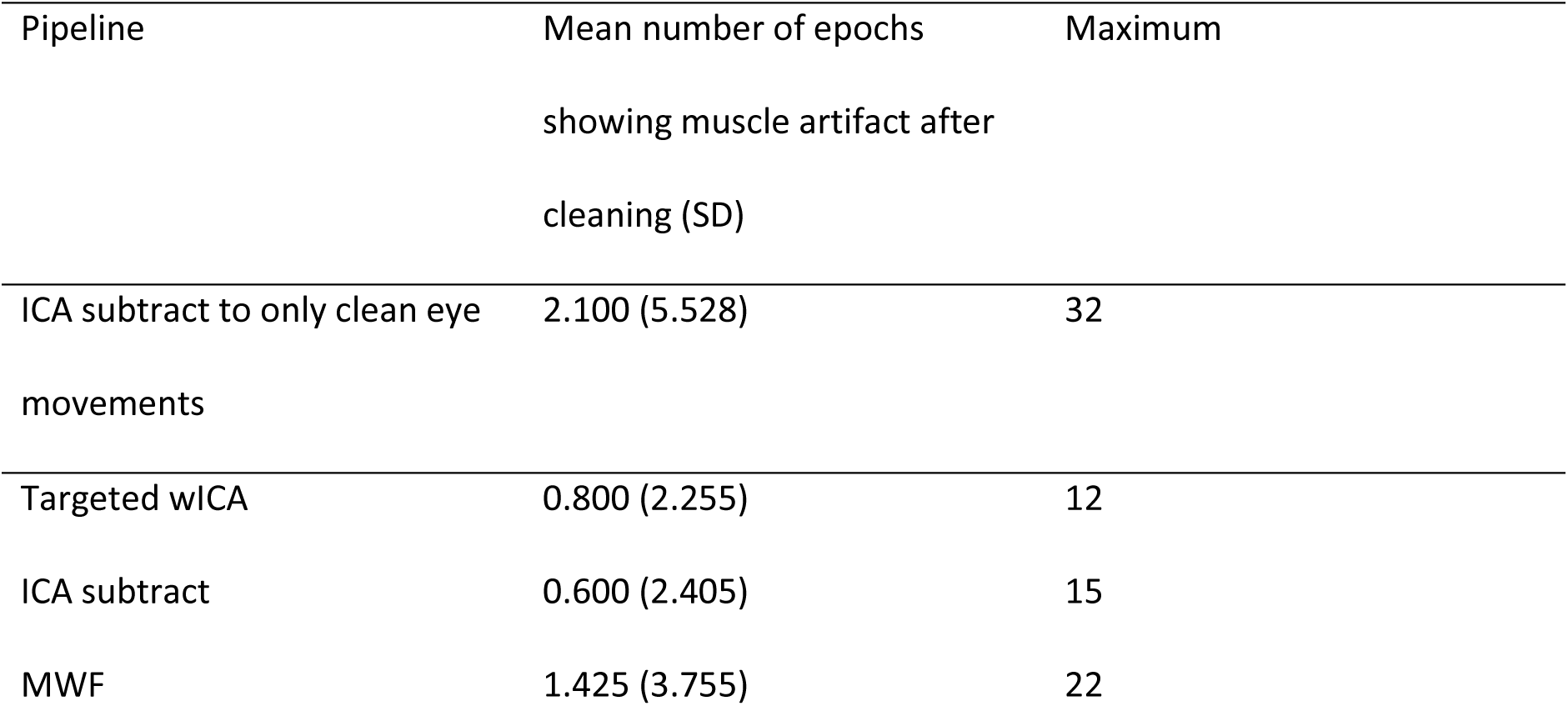
Mean number of epochs containing log-frequency log-power slopes indicative of muscle activity remaining in the N400 dataset after cleaning with each of the pipelines. ICA – independent component analysis, wICA – wavelet enhanced ICA, MWF – multi-channel Wiener filtering.

| Pipeline | Mean number of epochs<br>showing muscle artifact after<br>cleaning (SD) | Maximum |
| --- | --- | --- |
| ICA subtract to only clean eye<br>movements | 2.100 (5.528) | 32 |
| Targeted wICA | 0.800 (2.255) | 12 |
| ICA subtract | 0.600 (2.405) | 15 |
| MWF | 1.425 (3.755) | 22 |

### Effects of imperfect source component separation

#### ERP distortion

Next, our examination of the fronto-polar electrode ERPs from epochs that did not contain eye movement artifacts showed that all previously established methods that applied a component-based artifact reduction approach distorted the signal in epochs that did not contain eye movement artifacts. In particular, when we removed all non-eye-movement components (leaving data that characterised only the eye movement artifact), then reconstructed this eye-movement-artifact-only data into the scalp space, our data showed that ERP waveforms were captured within these artifact components, an effect that was particularly prominent at fronto-polar electrodes (Figures 7 and 8, left side). Critically, the ERP captured by the eye movement components differed between conditions for all pipelines, so when the artifact component was subtracted, the between condition difference was altered in these fronto-polar electrodes. This resulted in the between condition difference being inflated in the Go/No-go dataset and diminished in the N400 dataset (difference ERPs between the Go and No-go trials can be viewed in Figure 7, right side, and between the relevant and irrelevant trials of the N400 dataset in Figure 9, right side). Within the Go/No-go dataset, this led to larger ERP effect sizes for ERP comparisons that were conducted including all electrodes (Table 5). In contrast, targeted wICA was associated with only minimal distortion of the ERP constructed from epochs that did not contain eye movement artifacts. Targeted wICA also produced ERP difference topographies and effect sizes that were closely matched to the raw data from non-eye movement affected epochs.

**Figure 7.**
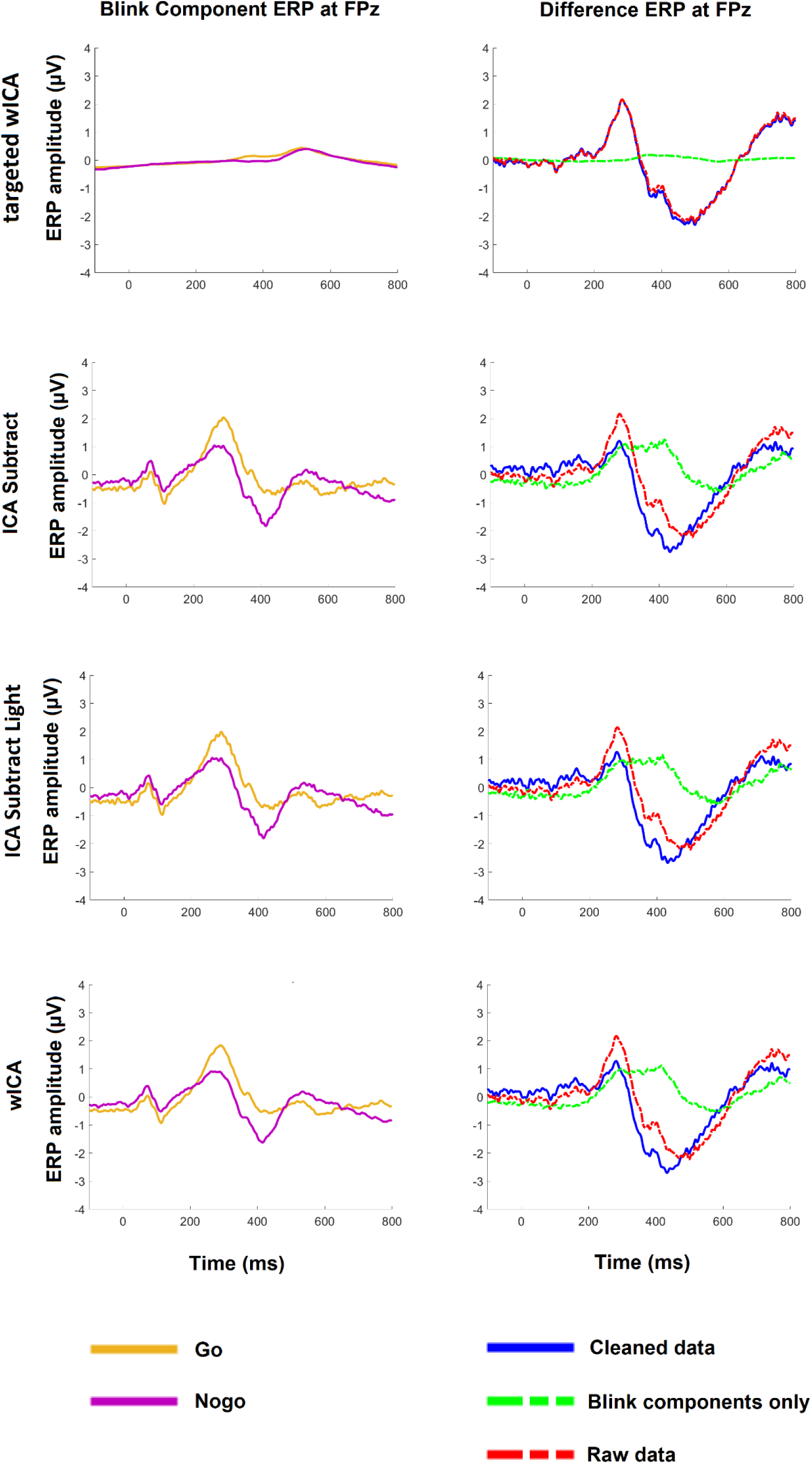
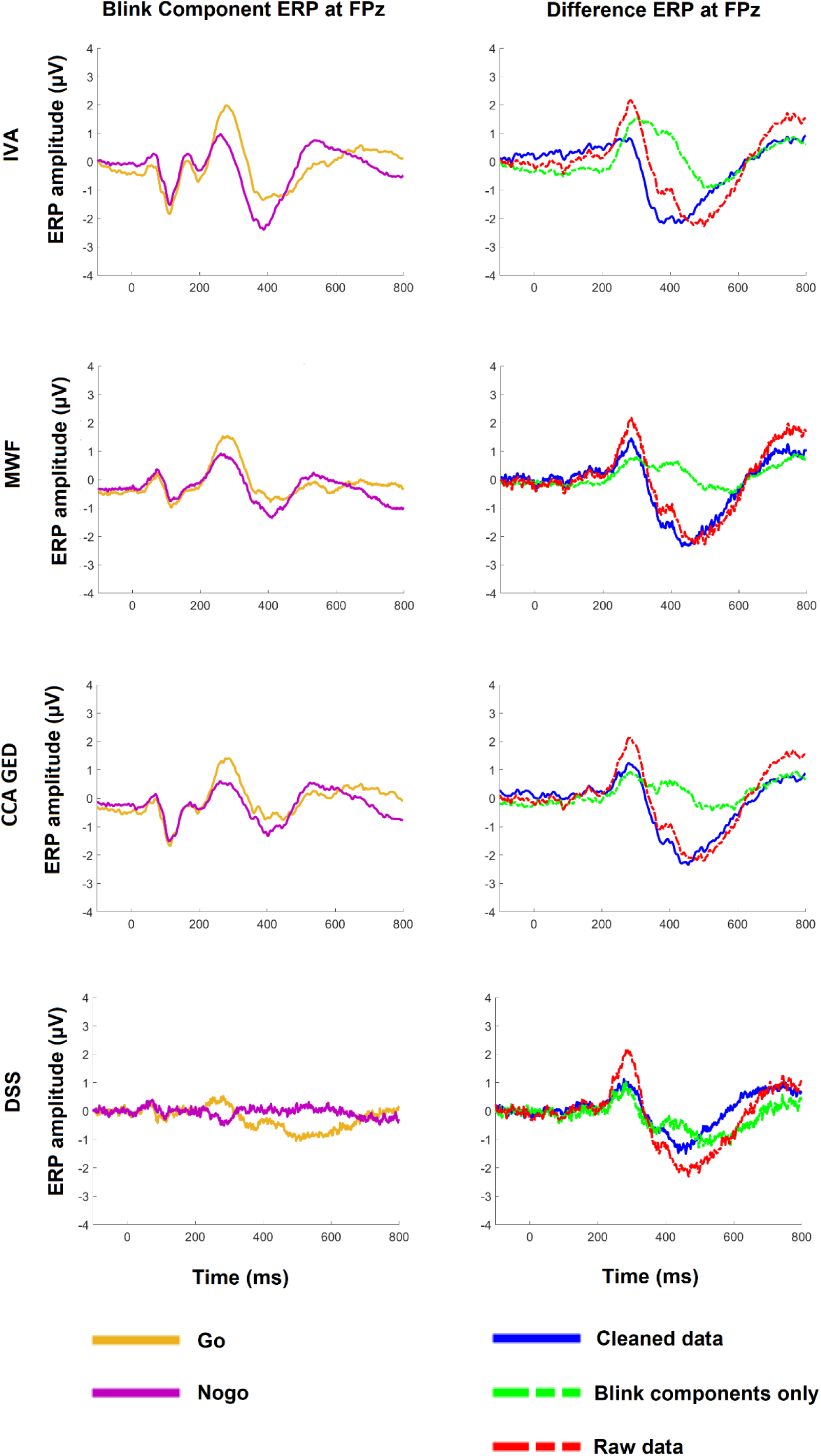
Left: the grand averaged event-related potential (ERP) signal at FPz obtained from the reconstructed eye movement artifact components during epochs that were not affected by eye movement artifacts for each of the different pipelines. Right: the effect that component-based reduction of this eye movement component had on the Go minus No-go difference ERP at FPz for each of the pipelines. Note that the MWF and DSS pipelines did not include the application of a low-pass filter, while all other data were low-pass filtered at 80Hz (as a result, the ERPs for these pipelines contain more high frequency activity than the ERPs from other pipelines). Note also that DSS cleans all inconsistent components simultaneously, so for the DSS pipeline we plotted the ERP within all artifact components rather than within only the eye movement components. ICA – independent component analysis, wICA – wavelet enhanced ICA, IVA – independent vector analysis, MWF – multi-channel Wiener filtering, CCA GED – canonical correlation analysis and generalised eigenvector decomposition, DSS – denoising source separation.

**Figure 8.**
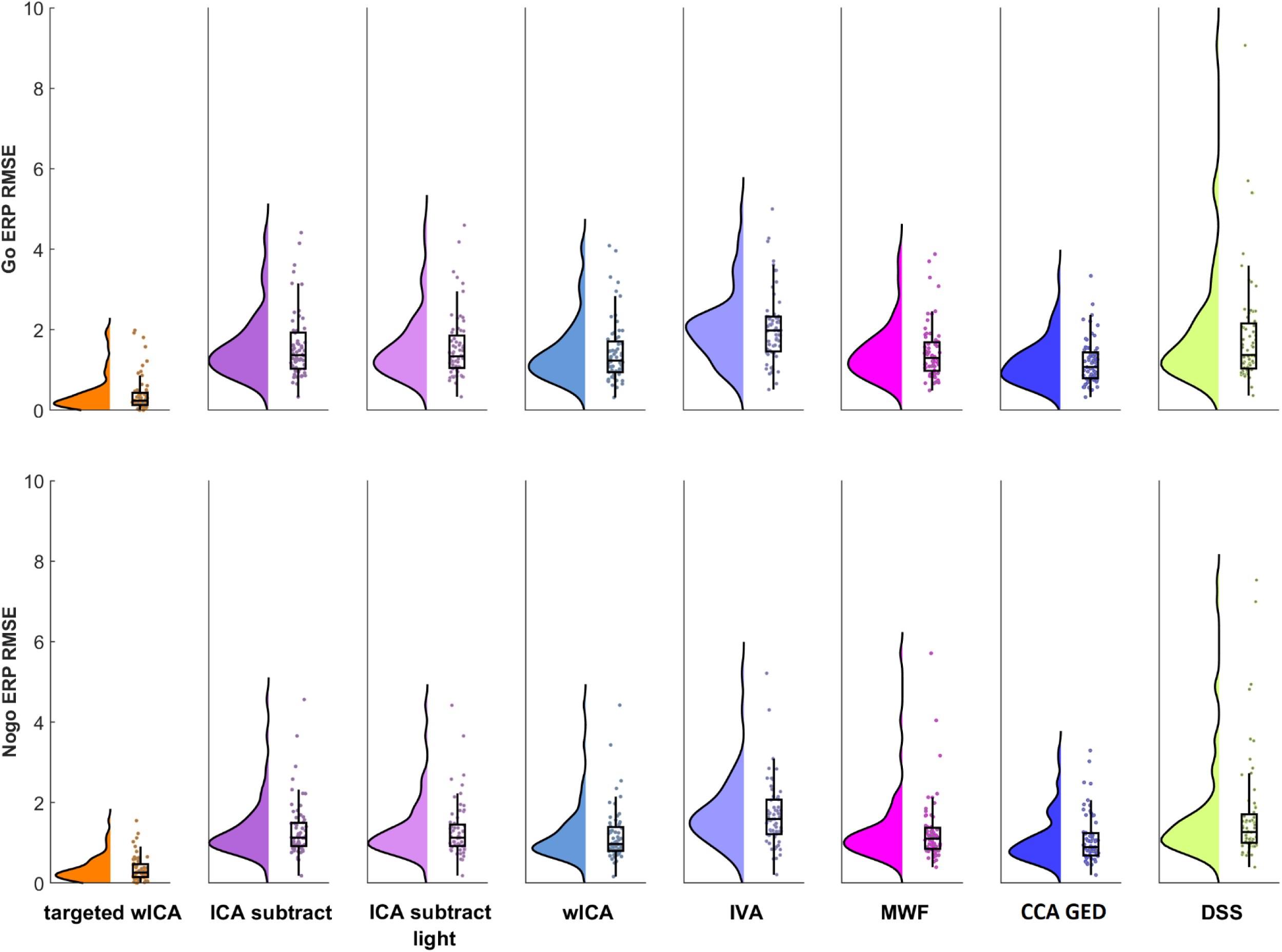
Root mean squared error (RMSE) values for each of the pipelines for the Go condition (above) and the No-go condition (below). Post-hoc t-tests indicated that the targeted wICA approach provided lower RMSE values than all other pipelines, indicating the least distortion of data outside of the artifact periods, while IVA provided higher RMSE values than all other pipelines (all pHolm < 0.05). No other pipelines showed significant differences to any other pipeline in the post-hoc t-tests. ICA – independent component analysis, wICA – wavelet enhanced ICA, IVA – independent vector analysis, MWF – multi-channel Wiener filtering, CCA GED – canonical correlation analysis and generalised eigenvector decomposition, DSS – denoising source separation.

**Figure 9.**
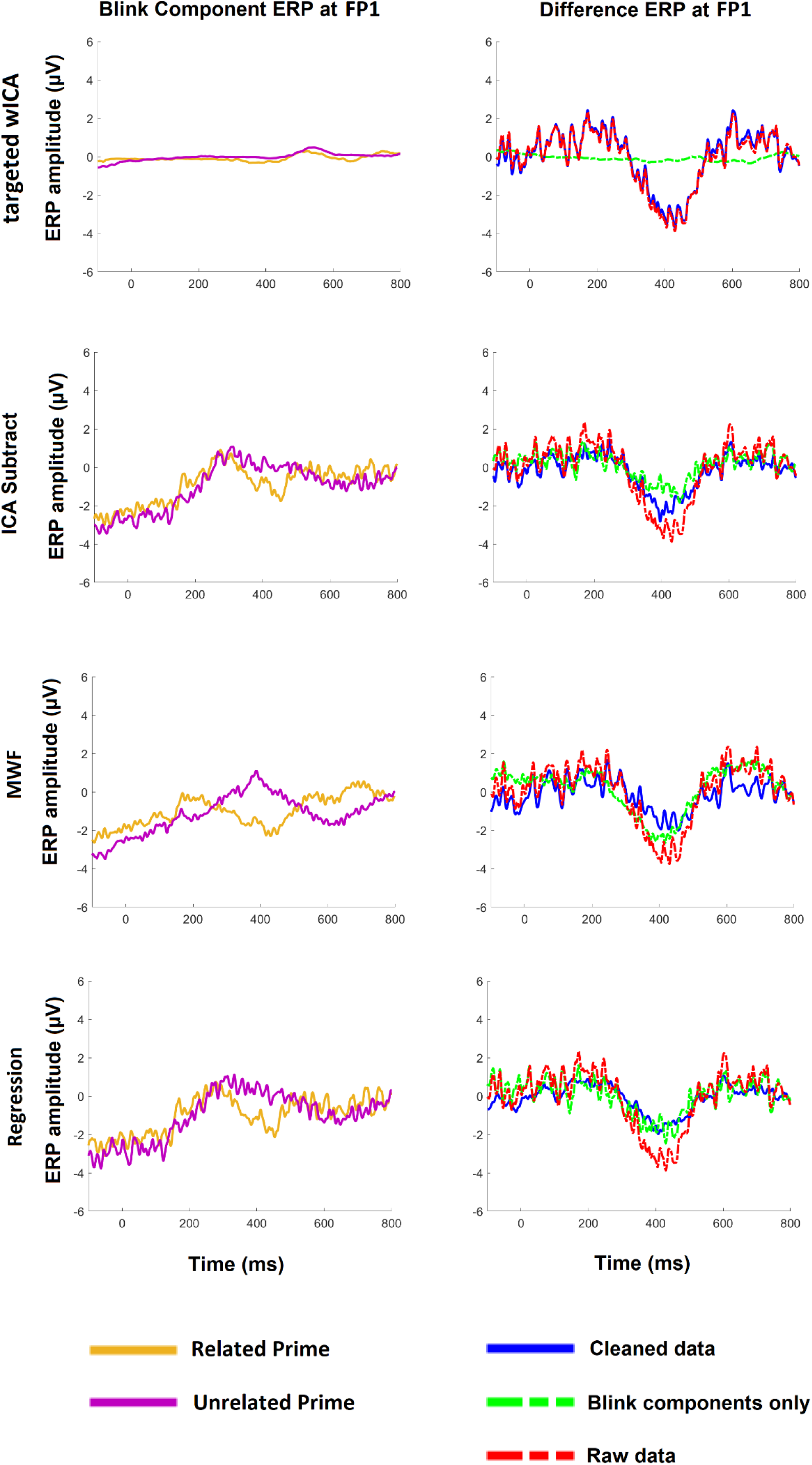
Left: the grand averaged event-related potential (ERP) signal at FP1 from the eye movement artifact components reconstructed into the scalp space during epochs that were not affected by eye movement artifacts for each of the different pipelines within the N400 dataset. Right: the effect that subtraction of this eye movement component had on the difference ERP for the related minus unrelated primes at FP1 for each of the pipelines. Note that when the eye-movement artifact component was reconstructed into the scalp space, the influence of this artifact component on fronto-polar electrodes contained a polarity inverted version of the N400. As such, eye movement correction removed some of the N400 activity from the FP1 electrode even in epochs that were unaffected by eye movement artifacts. In contrast, the targeted wICA approach avoided cleaning periods of the data that were not affected by eye movement artifacts, so it minimized the removal of the ERP activity that was mixed into eye movement components. ICA – independent component analysis, wICA – wavelet enhanced ICA, MWF – multi-channel Wiener filtering.

**Table 5.**
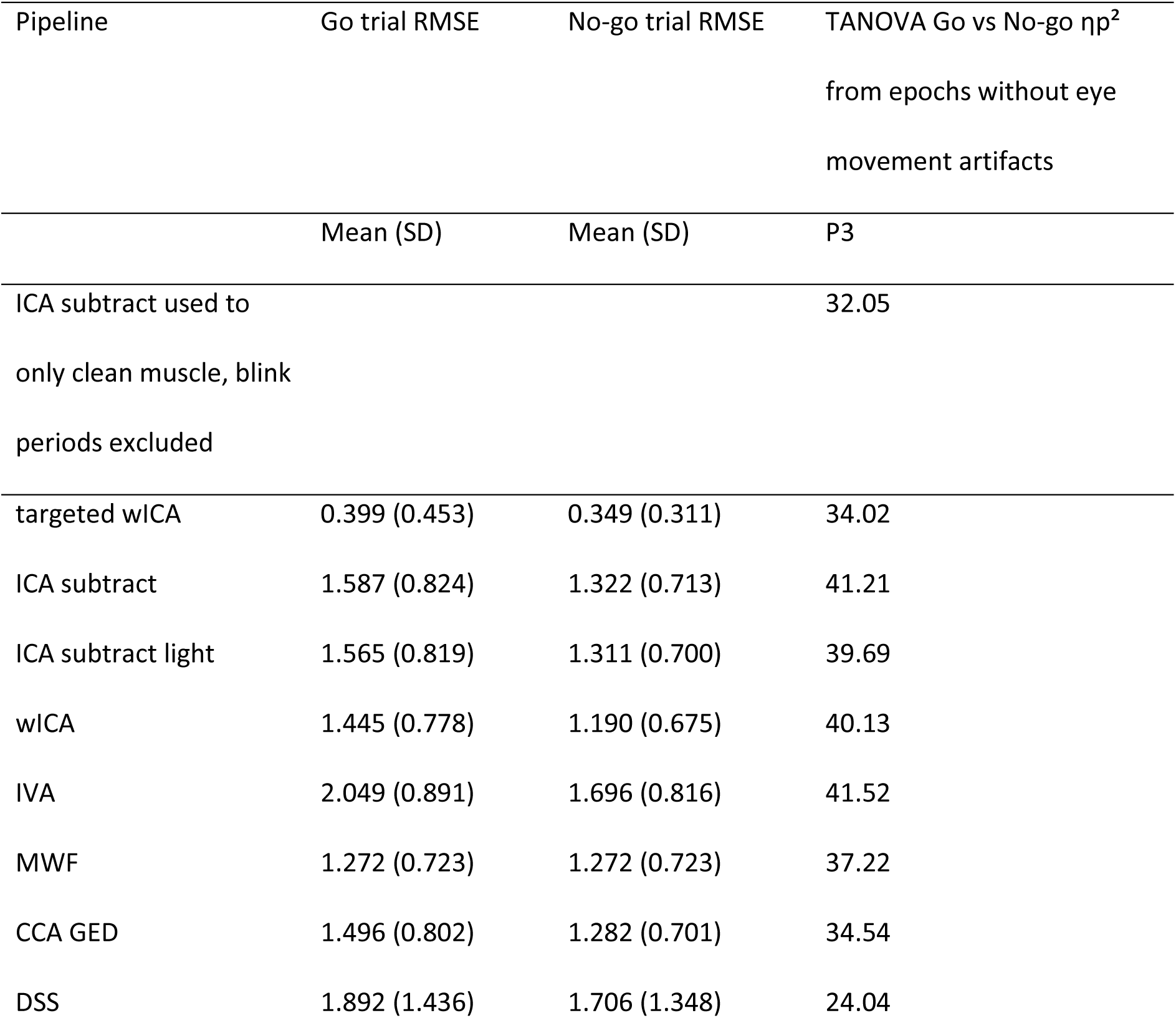
Root mean squared error (RMSE) values computed between ERPs at FPz constructed from raw data within periods that were not affected by eye movement artifacts and the ERP data after removing eye movement components. RMSE values are provided separately for Go and No-go trials. Effect sizes from TANOVA comparisons between Go and No-go trials after cleaning with each pipeline are also presented. Note that the targeted wICA method provided the lowest RMSE values, and TANOVA effect sizes that were matched most closely to the effect sizes from comparisons made after eye movement artifact components were left in the data and only epochs containing eye movements were excluded. ICA – independent component analysis, wICA – wavelet enhanced ICA, IVA – independent vector analysis, MWF – multi-channel Wiener filtering, CCA GED – canonical correlation analysis and generalised eigenvector decomposition, DSS – denoising source separation.

Figure 8 demonstrates the comparisons between pipelines of RMSE values computed between the cleaned data and the raw data. These comparisons were statistically significant within both the Go and No-go conditions (Go trials: F(7, 63) = 51.892, p < 0.001, ηp² = 0.452, ηG² = 0.220, No-go trials: F(7, 63) = 37.839, p < 0.001, ηp² = 0.375, ηG² = 0.198). Post-hoc t-tests indicated that across both Go and No-go trials, targeted wICA provided significantly lower RMSE values compared to all other pipelines (all pHolm < 0.001). For the Go trials, MWF also provided significantly lower RMSE values than ICA subtract, ICA subtract light, IVA, and DSS (all pHolm < 0.03), although MWF did not differ from these pipelines in the No-go trials (all pHolm > 0.90). In contrast, for both the Go and No-go trials, IVA and DSS performed worse than all other pipelines (pHolm < 0.001 for IVA and <0.021 for DSS) but did not differ from each other (pHolm = 0.748). wICA, ICA subtract, and CCA GED also did not differ from each other in either trial type (all pHolm > 0.60).

A similar effect was observed when muscle components were subtracted, where the ERP waveform was present in the scalp space reconstruction of the muscle components at the FT7 electrode - an electrode commonly affected by muscle activity (Supplementary Materials Figure S1, Section 3, page 15). However, while the muscle artifact components did contain this ERP waveform, the ERP waveform was small in amplitude (with a peak amplitude of less than 0.25µV) and did not differ between conditions, so component subtraction might not be predicted to inflate the between condition effect size.

Finally, the components that ICLabel identified as artifacts other than muscle and eye movement (for example, cardiac artifacts, channel and line noise artifacts, and unspecified artifacts) also contained ERP waveforms, demonstrating that these components identified as containing other artifacts also captured contribution from neural activity (Supplementary materials Figure S1, Section 3, page 15). However, the residual ERP captured within these non-eye-movement and non-muscle components was also small, with its maximal amplitude in the electrode space from grand average ERPs across all electrodes and timepoints being less than 0.25µV.

Similar results were apparent for the N400 dataset, with the ICA subtract, MWF, and the regression blink correction methods all producing distortion of the fronto-polar ERP waveform (Figure 9). However, in this case, the difference between conditions in the N400 ERP that was captured by the eye movement artifact component was in the same direction as the ERP effect, so the artifact cleaning resulted in a reduction in the size of the between condition effect at fronto-polar electrodes in the cleaned data. Again, the targeted wICA method produced the least distortion of the cleaned ERP.

The distortions produced by the ICA subtract, MWF, and regression pipelines resulted in significant differences in RMSE values computed between the cleaned data and the raw data within both the related and unrelated prime conditions (related trials: F(3, 26) = 39.351, p < 0.001, ηp² = 0.602, ηG² = 0.242, unrelated trials: F(3, 26) = 62.849, p < 0.001, ηp² = 0.707, ηG² = 0.378, Table 6). Post-hoc t-tests indicated that targeted wICA provided significantly lower RMSE values compared to all other pipelines (all pHolm < 0.001). The MWF pipeline performed better than ICA subtract and the regression pipelines (pHolm < 0.013 compared to ICA subtract, pHolm = 0.002 compared to the regression method). The ICA subtract and the regression pipelines did not differ from each other (pHolm = 0.446).

**Table 6.** Root mean squared error scores between ERPs constructed from raw data within periods that were not affected by eye movements and the ERP data after removing eye movement components for related and unrelated N400 trials. ICA – independent component analysis, wICA – wavelet enhanced ICA, MWF – multi-channel Wiener filtering.

| Pipeline | Related trial Mean | Unrelated trial Mean |
| --- | --- | --- |
|  | RMSE (SD) | RMSE (SD) |
| targeted wICA | 0.712 (1.036) | 0.386 (0.421) |
| ICA subtract | 2.753 (1.853) | 2.905 (1.721) |
| MWF | 2.326 (1.460) | 2.289 (1.286) |
| Regression | 3.001 (1.921) | 3.073 (1.723) |

#### Effects on connectivity

Our results also indicated that the component-based artifact reduction methods inflated dwPLI estimates of connectivity between FPz and Pz (Figure 10). After excluding epochs that contained eye movements, connectivity estimates from the cleaned data showed significant differences compared to the raw data. The ANOVA showed that this was the case both for Go trials: F(3, 63) = 30.304, p < 0.001, ηp² = 0.328, ηG² = 0.127, and No-go trials: F(3, 63) = 29.770, p < 0.001, ηp² = 0.324, ηG² = 0.122.

**Figure 10.**
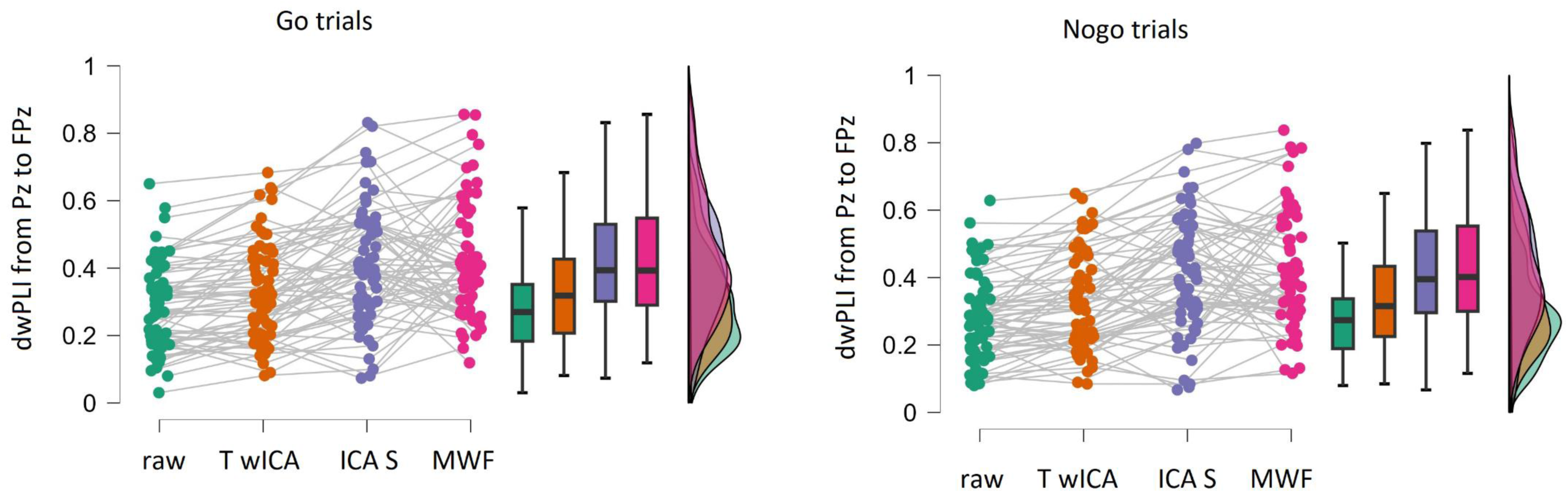
Debiased weighted phase lag index (dwPLI) values for the Pz to FPz electrode pairs from epochs that excluded eye movement artifacts for the raw data and after cleaning with each pipeline (T wICA = targeted wICA, ICA S = ICA subtract, MWF = multi-channel Wiener filtering). Statistical comparisons indicated that all pipelines showed higher dwPLI values than the raw data (all pHolm < 0.02), but also that the targeted wICA method showed less of a difference compared to the raw data in dWPLI values than the ICA subtract and MWF methods (all pHolm < 0.001).

Within the Go epochs, post-hoc tests indicated that all pipelines showed higher connectivity compared to the raw data (pHolm = 0.012 for the targeted wICA pipeline, and pHolm < 0.001 for all other pipelines). The ICA subtract and MWF pipelines also showed significantly higher connectivity than the targeted wICA pipeline (both pHolm < 0.001), but the ICA subtract and MWF pipelines did not significantly differ from each other (pHolm = 0.568). Within the No-go epochs, post-hoc tests showed that all pipelines showed higher connectivity values compared to the raw data (pHolm = 0.007 for the targeted wICA pipeline, and pHolm < 0.001 for all other pipelines). The ICA subtract and MWF pipelines also showed significantly higher connectivity than the targeted wICA pipeline (both pHolm< 0.001), but ICA subtract and MWF did not significantly differ from each other (pHolm = 0.499).

While the targeted wICA method reduced the inflation of connectivity estimates to provide a value that was closer to the value provided by the raw data, connectivity values after targeted wICA cleaning did still significantly differ from those provided by the raw data (see Figure 10). As such, our results indicate that using the targeted wICA cleaning is a good method to reduce the false positive connectivity that results from component subtraction artifact cleaning methods but it is not a perfect method. As such, if eye movement artifact periods can be excluded from the data without excluding too much of the data or adversely affecting analyses, excluding these periods may be preferable.

#### Effects on source localisation

In addition to the ERP and connectivity distortions created by component-based artifact reduction methods, source localisation of the primary generating dipole of the P3 produced dipole locations that were shifted more posterior relative to the ERP obtained from the raw data after excluding eye movement affected epochs (Figure 11). All pipelines provided a dipole located in approximately the left anterior cingulate cortex for No-go trials, and in the left thalamus for Go trials, in alignment with previous research showing dipoles in these regions during the Go/No-go task (DeLaRosa et al., 2020). However, the repeated measures ANOVA showed a significant difference between the component subtraction pipelines and the raw data in the Y location of the dipole (which reflects the anterior/posterior dipole location). For the Go P3: F(3, 63) = 13.284, p < 0.001, ηp² = 0.174, ηG² = 0.041, and for the No-go P3: F(3, 63) = 14.841, p < 0.001, ηp² = 0.191, ηG² = 0.047. Post-hoc tests indicated that for the Go P3, the Y location of the dipole from the ICA subtract and MWF pipelines was significantly more posterior than both the raw data (pHolm < 0.001), and the targeted wICA method (pHolm < 0.001). In contrast, the targeted wICA and raw data did not differ (pHolm = 0.772). The ICA subtract and MWF pipelines also did not differ from each other (pHolm = 0.772). Similarly, post-hoc tests for the No-go P3 indicated that the Y dipole for the ICA subtract and MWF pipelines was significantly more posterior compared to both the raw data (pHolm < 0.001) and the targeted wICA method (pHolm < 0.001). Again, the targeted wICA and raw data did not differ (pHolm = 0.831), and the ICA subtract and MWF pipelines also did not differ from each other (pHolm = 0.611).

**Figure 11.**
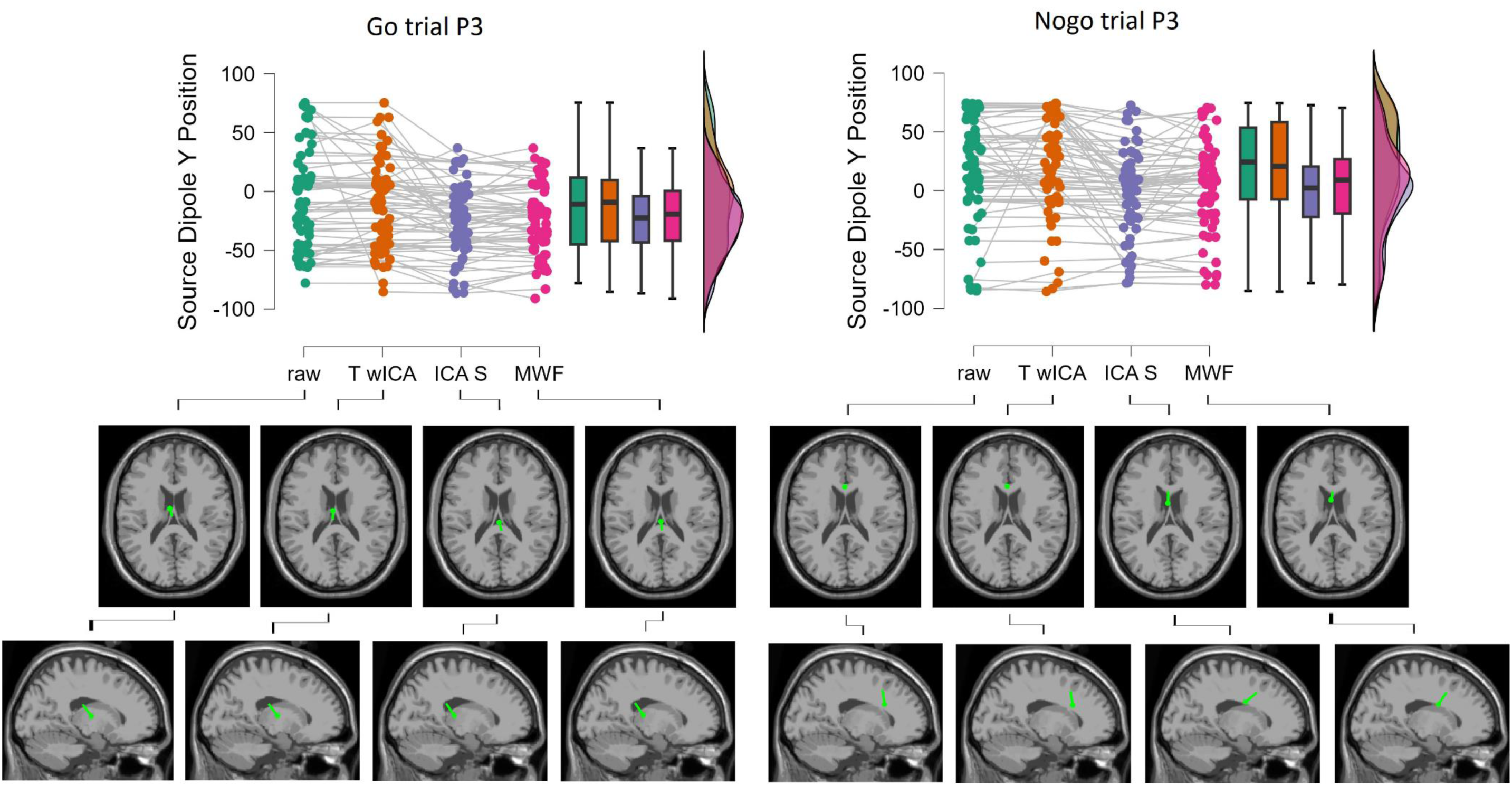
A raincloud plot depicting the source dipole Y position estimated for the Go and No-go P3 (top) and the mean location of the source dipole across all participants mapped onto the Montreal Neurological Imaging template MRI scan (bottom) after cleaning with four of the pipelines: raw data without any cleaning (but excluding eye movement affected epochs), our proposed targeted wavelet enhanced independent component analysis (T wICA) method, independent component analysis (ICA) subtract (ICA S), and multi-channel Wiener Filtering (MWF). Note that all ERP estimates were obtained after excluding epochs that contained probable eye movement artifacts. This ensured the P3 ERP was not impacted by the eye movement artifacts, so the results highlight the posterior shift in source localisation estimation resulting from removal of the eye movement components even in periods where no eye movement artifacts were present. The ICA subtract and MWF pipelines led to a minor but significantly more posterior P3 dipole estimation for both the Go and No-go trials compared to the raw data (pHolm < 0.001), while the dipole estimations after cleaning with targeted wICA did not differ from dipole estimations from the raw data (pHolm > 0.75).

## Discussion

A typical approach to address artifacts in EEG data is to decompose the data into component sources then subtract out components that are identified as representing artifacts, before reconstructing the scalp space data for analysis. Here, we show that because decomposition methods are imperfect, ERP waveforms are often mixed into artifact components, then removed from the data along with the artifact components. Counterintuitively, this can inflate experimental effect sizes, as the adverse effect of component-based artifact reduction can affect one condition more strongly than the other. In other cases, the adverse effects of component-based artifact reduction method can reduce effect sizes, in alignment with the common-sense prediction that reducing the neural signal will reduce effect sizes. Additionally, our results show that component-based artifact reduction methods can also inflate estimates of connectivity between electrodes and can shift the estimated source dipole location compared to analysing the raw data with eye movement artifact periods excluded.

Unfortunately, the adverse effect of imperfect component separation was ubiquitous across the component-based artifact reduction methods we tested, including ICA subtract, a less aggressive version of ICA subtract, IVA, CCA GED, and DSS. Furthermore, the issue was not resolved by methods that have been suggested to improve upon component subtraction approaches by taking temporal information into account, including wICA and MWF. Nor was it resolved by a regression eye movement correction method. To address this issue, we developed a novel approach that specifically targets artifact component cleaning to artifact periods (for eye movement artifact components) and frequencies (for muscle artifact components). Our results show that this targeted cleaning method provides artifact reduction that is equivalent or superior to previous methods, while also providing ERPs, connectivity estimates, and dipole localisation estimates that more closely match the raw data during non-artifact periods. As such, we recommend use of our targeted wICA cleaning method for a wide range of applications. To enable this, we have made a user-friendly version of our method available under an open access copyright within the RELAX EEG cleaning pipeline (which includes a graphical user interface to make the software user friendly). As such, the targeted wICA method is available through the RELAX plugin within EEGLAB, which is implemented within Matlab (https://github.com/NeilwBailey/RELAX).

To provide a full understanding of our results, we need to explain the counterintuitive finding that removing probable neural signal by subtracting an artifact component that contains that neural signal can inflate effect sizes. To provide this explanation, we first note that our results showed that the eye movement artifact component also contained some contribution from the No-go P3 ERP waveform, which component-based artifact reduction methods then reduced in the data (Figures 3 and 4). This No-go P3 is generated by fronto-central brain regions and produces a topographical pattern with a fronto-central positive maximum, where the opposite side of the generating source dipole is detected as a negative voltage shift at fronto-polar electrodes (Bekker, Kenemans, & Verbaten, 2005; Hong, Wang, Sun, Li, & Tong, 2017). In contrast, the Go P3 ERP has a topographical pattern with a more posterior maximum distribution (Bekker et al., 2005; Hong et al., 2017). We suggest that the proximity of the generating brain regions of the No-go P3 to the fronto-polar region increases the likelihood that or extent to which component-based methods will mix some contribution from the neural No-go P3 activity in with components that are identified as representing eye movement artifacts. This is likely due to an increased commonality between the No-go P3 and the eye movement artifacts in voltage shift patterns detected at frontal electrodes, in contrast to the more posteriorly distributed Go P3.

Since the No-go P3 ERP waveform in fronto-polar electrodes is characterised by negative voltages, removal of those negative voltages through component-based artifact rejection methods shifted the fronto-polar No-go P3 towards less negative voltages, increasing the difference at fronto-polar electrodes between the No-go P3 and the Go P3 (which was not as affected by the component rejection). In the raw data, the Go minus No-go difference was largest at the fronto-central electrodes, and the raw data showed only a medium sized difference between the Go and No-go P3 at fronto-polar electrodes (mean P3 amplitude difference = ∼1μV, Cohen’s d = 0.627). However, since the adverse effect of cleaning altered the amplitude of the fronto-polar No-go P3 more than the Go P3, the cleaned data showed a larger difference between the Go and No-go P3 at fronto-polar electrodes (mean P3 amplitude difference = ∼2μV, Cohen’s d = 1.051). In addition to affecting single electrode analyses, the alteration to the fronto-polar No-go ERP waveform also meant the difference topography of the Go minus No-go P3 was shifted to show more fronto-polar involvement, so effect sizes were also inflated in between condition comparisons that included all electrodes. In contrast, within the N400 dataset, the ERP waveform difference between conditions captured within the eye movement components showed the same direction of effect as the N400 effect present in the raw data. As a result, the component-based artifact reduction methods reduced the relevant vs irrelevant prime N400 effect size at fronto-polar electrodes instead of inflating the effect size.

In addition to the effects on ERP analyses, our results also indicated that component-based artifact reduction methods produced inflated estimates of connectivity between FPz and Pz, in alignment with previous research (Castellanos & Makarov, 2006). The explanation for this result is less obvious than for the inflated Go/No-go ERP effect size. In their paper introducing the wICA method, Castellanos and Makarov (2006) suggest that if subtracted components contain neural signals in addition to the artifact signal, then the subtraction of the neural signals from those components that are common to multiple electrodes will increase the commonality of signal shifts between those electrodes, inflating connectivity estimates. However, we note that the connectivity inflation they describe could only be the product of an increase in the commonality of instantaneous phase shifts between electrodes. The dwPLI method used in our study is robust against instantaneous phase shifts (as it is designed to be robust against spurious connectivity inflation due to volume conduction) (Vinck et al., 2011). As such, to explain the inflation of non-instantaneous phase connectivity effects in our study, we speculate that oscillations produced by fronto-polar sources would have been more likely to be removed by component-based artifact reduction methods. Due to the distance from the posterior sources, these fronto-polar sources may have contained phase shifts that were less likely to be synchronised to posterior sources, leading to low dwPLI estimates of the connectivity between FPz and Pz. However, it is likely that the signal at FPz also contained oscillatory activity generated by fronto-central sources (which may have reached FPz with a lower amplitude than the fronto-central electrodes due to the increased distance). The activity generated by these fronto-central brain regions may have contained phase shifts that were more strongly synchronised to posterior sources. If component subtraction removed more of the oscillatory signals that were generated by fronto-polar brain regions, the “signal to noise ratio” of oscillations generated by fronto-central regions may have increased, such that they had more prominent influence on the phase shifts in the signal recorded at FPz. As a result, the dwPLI connectivity estimates between FPz and Pz may have been increased by component-based artifact reduction methods, reflecting a spurious connectivity inflation generated by the removal of genuine neural signal. However, we note that this explanation is speculative, and future work is required to provide more definitive understanding of the issue.

Irrespective of the explanation for the increased connectivity after component-based artifact reduction, our results demonstrate that the common practice of using component-based artifact reduction methods is likely to lead to inflated estimates of functional connectivity in EEG research, with the potential for researchers to conclude functional associations are present between regions that are in fact not functionally associated. Although in our data connectivity was inflated for both the Go and No-go trials, we suggest this commonality of spurious connectivity inflation between experimental conditions might not be present for other experimental designs, so between condition effects might be spuriously inflated. Alternatively, in cases where a between condition difference is present, a spurious connectivity inflation in the condition with lower actual connectivity could mask the effect. Studies examining between group differences in connectivity might be vulnerable to spurious results if one group requires more components to be cleaned than the other (as might be the case when comparing psychiatric populations to healthy populations). In this situation, component-based artifact reduction could lead to a false positive difference in connectivity between the groups. Similar implications apply to studies using EEG to detect the effects of an intervention.

For example, a brain stimulation technique might be suggested to result in an increased engagement of frontal regions or an increase in connectivity between brain regions, when in fact the true result might be an increased activation of fronto-central brain regions that is spread more broadly by a component subtraction cleaning approach. While our proposed method did not completely resolve the issue of inflated connectivity estimates, it provided estimates that were significantly closer to the raw data (after excluding blink epochs) than our comparison methods. However, we note that if eye movement artifacts are infrequent and not aligned with task presentation, then perhaps it is better to simply exclude eye movement artifact periods from the connectivity estimation rather than risking inflating connectivity estimates by applying an artifact reduction method.

Finally, source dipole localisation uses the voltage shifts recorded at each electrode in combination with a function that models the putative underlying source volume conduction parameters to estimate the location of the brain regions generating each dipole (Oostendorp & Van Oosterom, 1989). Artifact cleaning methods like ICA subtract are commonly applied prior to source localisation, as source localisation estimates are sensitive to non-neural noise (Fatima, Quraan, Kovacevic, & McIntosh, 2013; Kaur, Singh, Bisht, Joshi, & Agrawal, 2022; Michel & Brunet, 2019; Niso et al., 2019; Stropahl, Bauer, Debener, & Bleichner, 2018). However, when the voltage shifts recorded at scalp electrodes are distorted by component-based artifact reduction, the source modelling algorithm receives biased data as its input. As a result, methods that rejected eye movement artifact components in the context of likely imperfect source separation produced source dipole location estimates that were significantly more posterior than those obtained from the raw data, even though all source estimates were computed from epochs that did not contain eye movement artifacts. This distortion has likely led to false conclusions about functional associations between cognitive processes and activity within specific brain regions in previous research.

Perhaps the application for which this issue is the most critical to resolve is in EEG source localisation for pre-surgical epilepsy evaluation, where source localisation methods are becomingly more commonly utilized (Van Mierlo, Vorderwülbecke, Staljanssens, Seeck, & Vulliémoz, 2020). Traditional component-based artifact rejection methods are commonly implemented to address artifacts prior to source localisation of the brain regions that generate focal epilepsies (Van Mierlo et al., 2020). Given our current results, this application of the ICA component rejection approach may lead to inaccurate estimation of the source of the epileptic activity. Our proposed pipeline addressed the issue of inaccurate source localisation for ERPs, providing dipole location estimates that did not differ from the estimates obtained from raw data after excluding blink periods. However, we note that in addition to improving the targeting of artifact reduction, future work might benefit from testing if it possible to develop methods that specifically filter seizure activities out of artifact components prior to component rejection, so that the seizure activity is preserved in the cleaned data.

It is worth noting that our results were demonstrated primarily with relevance to the cleaning of eye movement artifacts. In contrast, muscle components did not capture ERP activity that differed between the conditions, and only captured ERP activity of small amplitude (results reported in our Supplementary Materials, Figure S1, Section 3, page 15). However, because the muscle artifact components did contain at least some ERP contribution, it is possible that another experimental designs may produce signals that result in neural signals mixed into muscle artifact components containing a between condition difference. As such, we recommend future work refrains from applying component rejection methods to address muscle artifacts. Instead, we recommend muscle components are low-pass filtered at 15Hz (the approach provided by our targeted wICA method).

Similarly, the ERP waveform inadvertently mixed into the other artifact components (components identified as artifacts other than eye movements and muscles, i.e. cardiac, line and channel noise, and unspecified artifacts) was small, with a maximal grand average ERP amplitude across all single electrodes and timepoints being less than 0.25µV (results reported in our Supplementary Materials, Figure S1, Section 3, page 15). However, we note that the size of the averaged ERP mixed into these other artifact components might be higher in alternative experimental designs. Unfortunately, there is no obvious way to target these “other” artifacts, as they are not consistent and not as easily characterised as the blink and muscle artifacts. However, given that these other artifacts are not likely to be time-locked to stimuli (so are not likely to confound ERP analyses), it is likely prudent to avoid cleaning these artifacts, particularly if many epochs are available from which to construct the averaged ERP (which can remove random noise through averaging). If researchers do choose to clean these other artifacts, one potential method to address this could be to apply a wavelet transform to the artifact component in an attempt to extract only the artifact and preserve the ERP, as we have implemented in our previous research (Bailey, Hill, et al., 2023). However, our results demonstrate that this does not resolve the issue with regards to eye movement artifacts, and our informal testing suggests it is not resolved for other artifacts either. As such, we recommend against cleaning these non-eye-movement and non-muscle artifacts, and addressing potential issues caused by these artifacts by excluding affected epochs if necessary or by mitigating their effects through the grand averaging of a sufficient number of epochs. Overall, our results suggest that component-based artifact reduction methods alone are unlikely to sufficiently protect against distortion of the non-artifact data, even when enhanced by including temporal information as well as spatial information (as per the wICA, IVA, and MWF methods). As such, future attempts at progress on this issue might best focus on targeting cleaning at artifact periods or frequencies within artifact components, as exemplified by the targeted wICA method we introduce here, rather than simply focusing on attempting to improve the decomposition method. It may also be fruitful to identify methods that can specifically filter ERP-like data out of artifact components prior to rejection of those artifact components. Unfortunately, we were not able to identify a working method to achieve this, but future research may be able to solve this problem.

Our results also have implications for the testing methods used to validate future development of new EEG pre-processing pipelines. Most importantly, our results show that although it may seem desirable for a pipeline to maximise the detection of an effect of interest, an optimally performing pipeline must also be tested to confirm that it does not inflate a given effect size or project an effect to electrodes that previously did not show the effect. It is worth noting that our results also have implications for applications of component rejection approaches to clean data that extend beyond the field of EEG. In particular, we note that ICA is commonly used to clean fMRI data of motion, cardiac, respiratory, and scanner related artifacts (Bullock, Jackson, & Abbott, 2021).

In addition to our results regarding the effects of component-based artifact reduction methods, our preliminary tests of the optimal filter, electrode, and extreme artifact period rejection revealed some potentially important details (reported in full in the Supplementary Materials, Section 1). Interestingly, our tests of optimal filter settings indicates it may be worth questioning the conventional wisdom that <0.25Hz high-pass filter settings are optimal for ERP analyses, providing further support for a recent suggestion that 0.5Hz high-pass filtering may provide better results (Delorme, 2023). Between condition comparisons across all electrodes showed more variance explained by the experimental effect when high-pass filters were set at 0.5Hz to 1Hz. In particular, the P3 showed the largest effect size when high-pass filtered at 0.5Hz, the N400 at 0.75Hz, and the N2 at 1Hz. The full results are reported in our Supplementary materials, Section 1, pages 6-9, along with a discussion of relevant filtering issues. We also note that while our main results suggest cleaning methods can inflate effect sizes, it seems unlikely that the higher high-pass filter settings are inflating effect sizes. Filtering is achieved by multiplying each timepoint by a weighting transform function of the surrounding timepoints (de Cheveigné & Nelken, 2019). As such, it is unlikely (or perhaps even impossible) that a local maximum could be enhanced by filtering (and it is more likely for a local peak to be diminished), limiting the potential that filtering could inflate between condition effect sizes.

Additionally, both datasets showed larger effect sizes when bad electrodes and extreme artifact data periods were rejected using settings of light to moderate stringency prior to applying artifact reduction methods. This contrasts with our previously recommended RELAX default settings (which were higher in stringency / more aggressive, rejecting more electrodes and extreme artifact periods of the data). It is also in contrast to a recent approach suggested by (Delorme, 2023), where only a minimal number of electrodes are rejected and no extreme artifact periods were rejected. As such, it may be that ICA decomposition is still successful in the presence of moderately extreme outlying artifacts (so aggressive extreme artifact rejections do not need to be implemented) but ICA also does not perform as well when no extreme artifacts are addressed prior to ICA decomposition. This suggests a middle path is likely to be optimal for extreme artifact rejection settings. The full details of these analyses and a discussion about potential caveats can be found in our Supplementary Materials (Section 1).

### Limitations

While our results show the promising application of our novel targeted wICA method to address the issues produced by imperfect source decomposition, there are several limitations to our conclusions. First, we only tested two datasets (albeit medium to large datasets across different cognitive tasks, recording equipment and labs). As such, it remains a possibility that our results might not generalise to all other datasets. In particular, ICA decomposition is theoretically better with more electrodes (Klug & Gramann, 2021). It may be that recordings with >100 electrodes are not as affected by the issues associated with the component-based artifact reduction methods we report here, although we note that the suggested improvement with more electrodes may become practically negligible above 64 electrodes (Klug & Gramann, 2021). On a similar note, we tested for potential inaccurate source localisation using only 60 electrodes, a generic head model rather than individual MRI templates, and only examined the single dipole that explained the most variance. Higher electrode numbers and individual MRI templates are likely to increase source localisation accuracy (Michel & Brunet, 2019), and multiple dipole solutions are likely to more accurately characterise the P3 (Yamazaki et al., 2000). However, both our proposed targeted wICA method and the comparison methods were tested using the same methods, and we do not see an obvious reason why larger electrode numbers and individual MRI templates would address a posterior shift in source localisation produced by the ICA subtract and MWF methods, given the distortion is the result of signal distortion at the scalp prior to application of source localization.

Additionally, our tests only applied the PICARD ICA algorithm (Frank et al., 2022). However, our informal tests and the results of other researchers (Castellanos & Makarov, 2006; Chaumon et al., 2015) suggest that imperfect source separation is still an issue for other ICA algorithms, and recent research has suggested PICARD performs equivalently to the extended-infomax algorithm, which has been suggested to be the second best algorithm for EEG (second only to adaptive mixture ICA [AMICA]) (Frank et al., 2022). Furthermore, we tested IVA, GED, DSS and MWF methods, all of which showed the same issue as the ICA method. As such, we consider it unlikely that the issue could be resolved simply with the use of an alternative ICA algorithm (or even an alternative component rejection method).

Finally, targeted wICA provided blink amplitude ratios closer to 1 than other pipelines, indicating the absolute amplitude of activity within blink affect periods was equivalent to the absolute amplitude of activity outside of blink periods, suggesting more effective cleaning of blinks. However, we need to highlight that, in fact, the targeted wICA method reduced blink artifact amplitudes in exactly the same manner as the default wICA pipeline (with the exception that targeted wICA only affected periods of the data that were affected by eye-movement artifacts). Therefore, the superior blink amplitude ratio results for targeted wICA does not reflect superior blink cleaning. Instead, the advantage of targeted wICA within the blink amplitude ratio metric is that the amplitude of data outside of blink periods is not reduced by the wICA reduction of activity in eye movement components. As such, the blink amplitude results should be interpreted as showing that targeted wICA enables superior preservation of the neural signal outside of artifact periods, rather than superior blink artifact reduction.

### Conclusions

To summarise, we have provided evidence that the typically applied component rejection methods to address artifacts in EEG data can distort scalp-based ERPs, inflate connectivity estimates, and lead to inaccurate source localisation. Other commonly used artifact reduction methods do not resolve this issue. Our novel targeted wICA EEG cleaning approach addresses all these issues. The targeted wICA method produced ERPs, connectivity estimates, and source localisation estimates that were matched much more closely to raw data from non-blink epochs than all other artifact reduction approaches. Furthermore, our novel targeted wICA cleaning approach also addressed blink and muscle artifacts equivalently to or better than other cleaning methods, demonstrating its promise for both preserving signal and effectively removing artifacts. To enable application of our targeted wICA method, we have provided the code as part of a freely available and easy-to-use plugin for EEGLAB: the RELAX open access software (https://github.com/NeilwBailey/RELAX).

## Funding Information

PBF is supported by a National Health and Medical Research Council of Australia Practitioner Fellowship (6069070). No funding was provided specifically for this project.

## Conflict of Interest

In the last 3 years PBF has received equipment for research from Neurosoft, Nexstim and Brainsway Ltd. He has served on scientific advisory boards for Magstim and LivaNova and received speaker fees from Otsuka. He has also acted as a founder and board member for TMS Clinics Australia and Resonance Therapeutics. PBF is supported by a National Health and Medical Research Council of Australia Investigator grant (1193596). The other authors declare that they have no conflicts of interest.

## Ethics Information

Ethics approval for the first dataset was provided by the Ethics Committees of Monash University and Alfred Hospital, and ethics approval for the second dataset was provided by the Institutional Review Board at the University of California, Davis. All participants provided written informed consent prior to participation in the study.

## Data Availability Statement

The data that support the findings of this study are available from the corresponding author, NWB, upon reasonable request, and from the ERPCore dataset.

## Code availability

https://github.com/NeilwBailey/RELAX

