## Supplementary Materials for "EEG is better when cleaning effectively targets artifacts"

***Section 1: Testing of Initial Pre-processing Parameter Settings - Methods***

Prior to our primary analyses, in order to determine which aspects of the data our primary analyses should focus on, we first examined the ERP data without any artifact cleaning to determine the ERP periods that showed potential experimental effects that seemed worth examining in our primary analyses. To achieve this, we tested the effect sizes produced by between condition comparisons of the data without the application of any artifact reduction method, but instead after the exclusion of extreme artifact data segments, electrodes, and periods affected by blinks after bandpass filtering the data (using a range of bandpass limits, described below). While we note that a primary message of our study is that the optimization of experimental effect sizes can be a misleading metric to assess optimal data cleaning due to the potential inflation of effect sizes by component-based artifact reduction methods, the issue is unlikely to apply when assessing effect sizes without implementing any component reduction method. As such, assessment of the effects of different filtering approaches and different extreme artifact rejection settings on effect sizes without the application of a component subtraction artifact reduction method is still a valid method to determine optimal settings. To determine effect sizes, we used a topographical ANOVA (TANOVA) (Koenig, Kottlow, Stein, & Melie-García, 2011). The TANOVA subtracts the values at all electrodes in the first condition from the mean at all electrodes from the second condition, followed by the calculation of the global field potential on this difference topography, which produces a global dissimilarity topography that reflects the overall difference between conditions including all electrodes after cleaning by each pipeline (Koenig et al., 2011).

Within the Go/No-go data, there were significant and prominent between condition effects within the N2 window from 185 to 315ms and the P3 window from 315 to 500ms. Within the N400 data, there were significant and prominent between condition effect from 340 to 480ms. These time windows align with time windows for these ERPs used in previous research. As such, these periods were used to test the effects of different filtering, extreme artifact rejection, and artifact cleaning approaches in our primary analyses. We note that the between condition effects were statistically significant for longer durations after artifacts were removed (in alignment with the intended goal of artifact reduction methods). Nonetheless, for our primary analyses we restricted our comparisons to the time-window that showed significant between condition differences prior to artifact cleaning. This ensured our primary comparisons were not biased by any particular artifact cleaning method, protecting our results against potential biases that might be caused by a scenario where a specific artifact reduction approach prolonged between condition ERP effects while other artifact reduction approaches did not.

Next, in addition to the primary analyses reported in the main manuscript, we explored the pre-processing settings that preceded the component-based artifact reduction. These pre-processing settings included commonly applied bandpass filtering, as well as EEG electrode and period rejection steps based on extreme outlying artifacts. Prior to applying artifact reduction methods, it is typical to filter the data and reject electrodes and data periods that are extremely contaminated with artifacts (under the assumption that any signals lying under the extreme artifact signals are likely to be irretrievable, or that the artifact is so severe its inclusion will adversely affect the performance of component-based artifact reduction approaches). Filtering the data reduces the contribution of extremely high frequency activity that is more likely to reflect muscle activity or artifacts than brain activity (Muthukumaraswamy, 2013) and extremely low frequency activity that is unlikely to contain a neural response to stimuli (Rousselet, 2012). High-pass filtering approaches are commonly implemented to reduce the confounding effects of very low frequency drift on ERPs (Rousselet, 2012). However, filter settings are debated. Traditional perspectives suggest that setting high-pass filters to exclude frequencies above 0.25Hz can reduce the amplitude of late ERPs (Rousselet, 2012; Tanner, Morgan-Short, & Luck, 2015). In contrast, recent research has suggested that high-pass settings of 0.5Hz or higher can more optimally detect effects of interest (Delorme, 2023). Furthermore, ICA performance is adversely affected when frequencies below 1Hz are included

(Winkler, Debener, Müller, & Tangermann, 2015), and we have demonstrated a similar performance reduction for MWF cleaning approaches (Bailey, Hill, et al., 2023). Furthermore, because filtering the data involves performing a mathematical transform on each timepoint based on the surrounding timepoints, filtering can introduce its own artifact contribution to the data, particularly when large amplitude spikes are present. As such, robust detrending has been suggested as an alternative to filtering the data (de Cheveigné & Arzounian, 2018).

In an attempt to determine optimal filter settings for our data (which we could use in our primary analyses), we tested the effects of applying fourth order acausal Butterworth high-pass filters at 0.25Hz, 0.5Hz, 0.75Hz and 1Hz to the ERP comparisons without any artifact reduction methods applied. We also tested the application of a second order polynomial robust detrending approach instead of any high-pass filter (de Cheveigné & Nelken, 2019). Additionally, to ensure the results of these tests were not simply related to artifacts remaining in the data, and to ensure the filter or robust detrending settings did not interact with the artifact cleaning approaches, we tested effect sizes provided by the same settings followed by artifact reduction using the wICA default cleaning within RELAX (Bailey, Hill, et al., 2023) for the Go Nogo dataset, and using our newly developed targeted wICA cleaning for the N400 dataset. The default wICA cleaning was applied to the Go Nogo dataset as our first test of potential interactions between filtering and artifact cleaning pipelines as a reasonable default cleaning approach to ensure our selection of filter settings was not biased towards a specific cleaning approach. However, after we had established filter settings with these initial tests, we used targeted wICA to test for potential interactions between filtering and cleaning within the N400 dataset. For these post-artifact reduction comparisons, we included all epochs, under the assumption that artifacts were effectively cleaned by the artifact reduction method. We note that these comparisons may be influenced by the effect size inflation that can be produced by artifact component reduction approaches. As such, we interpreted these results with caution, and by referring to the results that were produced without applying any artifact reduction to check for alignment between the results following artifact reduction and the results that were robust to potential effect size inflation.

Similar to the debate around filter settings, there is no agreed consensus on how contaminated by artifacts an EEG electrode or data period from the raw data should be before it is entirely rejected prior to any further artifact reduction. Our perspective has been that data periods that are so contaminated they are unlikely to contain retrievable neural data should be rejected, and electrodes that are contaminated by artifacts for a duration that is extensive enough that their inclusion would likely adversely affect ICA decompositions or lead to rejection of an excessive number of epochs should be rejected (Bailey, Biabani, et al., 2023). However, recent work has suggested that minimal electrode rejection and no rejection of data periods is an optimal approach (Delorme, 2023).

To address this uncertainty, we tested the effect of different extreme outlying data exclusion settings prior to artifact reduction methods, on outcomes of between condition comparisons, and on the degree of ERP distortion produced by the artifact reduction methods applied after the extreme artifact rejection step. We tested four different approaches, which are summarised in Supplementary Table S1: 1) the minimal approach suggested by Delorme (2023), where electrodes that showed a correlation below $r < 0.9$ with any other electrode were excluded, as were electrodes that showed 50 or 60Hz line noise that was more than 4SD from the mean of all electrodes, but no data period exclusions were implemented; 2) the RELAX default electrode and extreme period rejections (which are reasonably aggressive). This approach first selects electrodes for deletion based on which electrodes are most severely contaminated by artifacts (with a maximum of 20% of electrodes able to be rejected). It then selects the worst data periods for rejection using the following methods: outlier detection within the distribution of voltage values (>8 SD from the mean), the kurtosis of voltage values (>8 SD from the mean), extreme voltage values (>500µV), extreme voltage shifts, and extreme log-power log-frequency slopes reflecting either high amplitude low frequency drift or very large amplitude high-frequency muscle activity within each 1 second period of the data; 3) a more moderate stringency RELAX electrode and extreme period rejection approach, which involved less aggressive settings so fewer electrodes and data periods were rejected. This moderate approach allowed a maximum of 10% of electrodes able to be rejected, and set extreme rejection thresholds to >10 SD from the mean, and extreme voltage thresholds to > 1000µV; and 4) A light version of the RELAX electrode and extreme period rejection approach, which allowed a maximum of 10% of electrodes able to be rejected, and with minimally aggressive settings so even less data were rejected. The thresholds for rejections under this version were set to >12 SD from the mean and the extreme voltage threshold was set to > 1000µV.

| Parameter | RELAX default (stringent) | RELAX moderate | RELAX light | Minimal (Delorme) |
|---|---|---|---|---|
| Maximum voltage shift within each 1s period | 20 MAD from the median of all epochs | 25 MAD from the median of all epochs | 30 MAD from the median of all epochs | N/A |
| Maximum voltage shift within each blink affected period | 8 MAD from the median of all epochs | 10 MAD from the median of all blink affected periods | 12 MAD from the median of all blink affected periods | N/A |
| Absolute voltage threshold | 500µV | 1000µV | 1500µV | N/A |
| Improbable voltage distribution | 8SD | 10SD | 12SD | N/A |
| Kurtosis | 8SD | 10SD | 12SD | N/A |
| Proportion of time contaminated by muscle before an electrode can be rejected (log-power log-frequency slope for detecting muscle) | 0.05 (-0.59) | 0.50 (-0.31) | 0.50 (-0.31) | N/A |
| Proportion of time contaminated by extreme artifacts before an electrode is rejected | 0.05 | 0.25 | 0.40 | N/A |

| | | | | |
|---|---|---|---|---|
| Max proportion of electrodes that can be removed | 0.20 | 0.10 | 0.10 | N/A |
| Line noise electrode rejection threshold | N/A | N/A | N/A | 4SD |
| Correlation with other electrodes required | N/A | N/A | N/A | 0.9 |

**Supplementary Table S1.** Extremely bad electrode and EEG data period rejection settings that were tested in our preliminary tests to determine the optimal settings for our primary analyses. MAD = median absolute deviation.

***Testing of Initial Pre-processing Parameter Settings - Results***

Overall, results indicated that high-pass filtering the data at 0.5Hz was optimal for detecting differences in the Go/Nogo P3, while high-pass filtering at 1Hz was optimal for the Go/Nogo N2 (Table S2). These settings were optimal both when the data were tested without any artifact reduction and when the data were cleaned with wICA, suggesting the effect of filter settings did not interact with the wICA artifact reduction method. However, within the N400 dataset, data without any artifact reduction applied suggested that high-pass filtering at 0.5Hz filtering was best, while if targeted wICA was used to reduce artifacts in the data, high-pass filtering at 0.75Hz performed best. To ensure the results of our primary tests were not biased by an interaction between the high-pass filter settings and the cleaning method, and in an attempt to minimize the potential adverse effects of filtering, we high-pass filtered all data at 0.5Hz prior to our primary comparisons. This is also in alignment with the optimal settings for the data before any artifact reduction method was applied for both the P3 and N400. However, we note that if artifact reduction methods are applied in future research, then high-pass filtering at 0.75Hz might be more effective for the N400. We also note that future research may find high-pass filtering at 1Hz is more optimal for examining the N2.

We note that these results conflict with previous suggestion that high-pass filtering settings above 0.25Hz adversely affect analysis of the P3 (Rousselet, 2012; Tanner et al., 2015). High-pass filtering above 0.3Hz has been suggested to reduce late-latency ERP amplitude and produce filter artifacts such that an ERP peak can be surrounded by a filter-artifact effect that shows an inverted polarity ERP (Tanner et al., 2015). This would argue against the application of high-pass filter settings at 0.5Hz to 1Hz. However, we note that despite the potential presence of the filter artifacts in our data, the maximum effect sizes for the N2, N400, and P3 at a single timepoint (at the ERP's maximum) were larger when our data were high-pass filtered at 1Hz, 0.75Hz, and 0.5Hz respectively. While the inverted polarity filter artifacts demonstrated by Tanner et al. (2015) could produce an inflated effect size on either side of an ERP's peak, we have not seen any evidence that a filter artifact could enhance an effect size at the ERP's peak, or adversely affect the ERP waveform for only a single condition and not another condition. Indeed, since filtering is achieved by multiplying each timepoint by a weighting transform function of the surrounding timepoints (de Cheveigné & Nelken, 2019), it is unlikely (or perhaps even impossible) that a local maximum could be enhanced by filtering (and it is more likely for a local peak to be diminished). This suggests that although filtering artifacts may be an issue, it seems likely that at least in the two datasets we tested, the presence of low frequency drift that reduced the ERP effect was more of an issue, and that high-pass filter settings are optimal at 0.5Hz or higher. Furthermore, although the application of filters to EEG data has been suggested to be non-optimal (de Cheveigné & Nelken, 2019), high-pass filtering the data was associated with larger effect sizes for all datasets and ERPs than the robust detrending approach suggested by de Cheveigné and Arzounian (2018).

One potential reason for our filtering result is that it may be possible that high-pass filter settings above 0.25Hz do reduce late ERP amplitudes, but also mitigate drift confounds that reduce detection of the between condition effects of interest. In particular, we note that our Go/No-go task displayed stimuli at 0.9Hz with a random 50ms jitter which commonly produced a low frequency oscillation that was synchronised to the stimulus presentation timing. Additionally, our data often showed prominent low frequency drift. Reducing the low frequency drift and slow oscillation synchronised to stimuli by using higher high-pass filter settings may have improved the ability to distinguish between the two conditions, despite potentially reducing the P3 amplitude. Delorme (2023) similarly reported improved performance with higher high-pass settings (>0.5Hz), suggesting that our findings are not an anomaly and perhaps conventional wisdom on high-pass filtering settings should be reconsidered. Given that our results indicated 0.5Hz high-pass filtering produced optimal effect sizes when no artifact reduction methods were applied for two out of the three ERPs, we applied the 0.5Hz high-pass filtering in our primary analyses. This is with the exception of the DSS method, where we replicated the approach suggested by de Cheveigne (2023) as closely as possible (including applying robust detrending rather than filtering prior to cleaning).

Next, results indicated that applying extreme electrode and data period rejections with the moderate stringency or light stringency settings using RELAX provided larger effect sizes than other extreme rejection settings, depending on the dataset and ERP analysed (Table S3). Specifically, moderate stringency extreme rejections led to more variance explained for the Go Nogo P3, while light stringency settings led to (slightly) more variance explained for the Go Nogo N2, and more variance explained for the N400. We also tested the amount of ERP distortion that resulted from different extreme artifact rejection settings prior to applying a default wICA cleaning (for the Go/No-go dataset) and prior to applying targeted wICA (for the N400 dataset). As per our primary comparisons, we achieved this by computing the RMSE between the ERP after extreme artifact rejections but prior to application of the wICA artifact reduction method to the ERP after both the same extreme artifact rejections and after wICA cleaning, with all ERPs obtained from a fronto-polar electrode (FPz in the Go/No-go data and FP1 in the N400 data). However, in contrast to our primary results (which report the RMSE values after only reducing eye movement artifact components), for these analyses, to provide a test that aligns with the real world application of artifact reduction methods, we used wICA to clean all types of artifacts (using RELAX's wICA default for the Go/No-go dataset), or both eye movement and muscle artifacts (using targeted wICA for the N400 dataset). We note that including these other artifacts in the cleaning approach led to higher RMSE values than was the case for our primary analyses. These comparisons showed a significant difference between the different extreme artifact rejection settings for the N400 dataset: $F(3, 26) = 3.476$, $p = 0.030$, $\eta p^2 = 0.118$, $\eta G^2 = 0.032$ (see Table S4). Post-hoc t-tests within this N400 dataset indicated that the effect was driven by the RELAX moderate extreme artifact rejection settings performing better than the extreme artifact rejection settings proposed by Delorme (2023) (pHolm = 0.046), while other pairwise comparisons did not significantly differ (all pHolm > 0.010). However, there was no significant difference between the different extreme artifact rejection settings for the Go/No-go dataset, where RELAX's wICA default approach was used to reduce artifactual components: $F(3,63) = 2.295$, $p = 0.115$, $\eta p^2 = 0.035$, $\eta G^2 = 0.002$.

| **High-pass filter settings** | **Go Nogo N2** | **Go Nogo P3** | **Relevant/Irrelevant N400** |
|---|---|---|---|
| **No cleaning** | | | |
| Robust detrending | 21.98 | 26.37 | 7.65 |
| 0.25Hz | 22.36 | 24.51 | 15.49 |
| 0.5Hz | 24.71 | **30.07** | **18.67** |
| 0.75Hz | 25.71 | 30.01 | 17.92 |
| 1Hz | **29.42** | 29.49 | 18.21 |
| **wICA** | | | |
| Robust detrending | 22.55 | 34.35 | 26.20 |
| 0.25Hz | 25.39 | 43.61 | 25.92 |
| 0.5Hz | 25.51 | **43.78** | 32.77 |
| 0.75Hz | 33.93 | 43.53 | **34.87** |
| 1Hz | **36.09** | 41.93 | 32.79 |

**Supplementary Table S2.** $\eta p^2$ effect sizes for TANOVA comparisons between Go and Nogo trials using different filter settings, followed by the exclusion of eye movement affected epochs and grand averaging to obtain ERPs (above), or the reduction of artifacts using wICA (for the Go/Nogo dataset, below) or targeted wICA (for the N400 dataset) and grant averaging to obtain ERPs from all epochs (including eye movement artifact epochs, as these were reduced by wICA cleaning).

| **Extreme electrode and data period rejections** | **Go Nogo N2** | **Go Nogo P3** | **Relevant/Irrelevant N400** |
|---|---|---|---|
| **0.5Hz high-pass filter** | | | |

| | | | |
|---|---|---|---|
| RELAX defaults (high stringency) | 25.51 | 43.78 | 29.36 |
| RELAX moderate stringency | 29.31 | **44.41** | 32.77 |
| RELAX light stringency | **29.48** | 43.88 | **35.48** |
| Minimal rejections (Delorme) | 27.73 | 42.76 | 29.15 |
| **1Hz high-pass filter** | | | |
| RELAX defaults (high stringency) | 36.09 | 41.93 | These tests were not performed, as the 1Hz high pass filter setting was not optimal for the N400 ERP. |
| RELAX moderate stringency | 35.83 | 41.95 | |
| RELAX light stringency | **36.14** | **42.34** | |
| Minimal rejections (Delorme) | 24.86 | 34.47 | |

**Supplementary Table S3.** $\eta p^2$ effect sizes for TANOVA comparisons between conditions after applying different extreme artifact rejection settings, followed by the reduction of artifacts using wICA (for the Go/Nogo dataset, above) or targeted wICA (for the N400 dataset, below) and grant averaging to obtain ERPs from all epochs (including blink affected epochs, as these were reduced by artifact cleaning).

| **Extreme electrode and data period rejections** | **Go/No-go**<br>**Go Mean (SD)**<br>**Nogo Mean (SD)** | **Relevant/Irrelevant**<br>**Relevant Mean (SD)**<br>**Irrelevant Mean (SD)** |
|---|---|---|
| **0.5Hz high-pass filter** | | |

| | | |
|---|---|---|
| RELAX defaults (high stringency) | 1.598 (0.745)<br><br>1.286 (0.623) | 0.868 (1.098)<br><br>0.776 (0.817) |
| RELAX moderate stringency | 1.607 (0.731)<br><br>1.286 (0.594) | **0.774 (0.774)**<br><br>**0.726 (0.789)** |
| RELAX light stringency | 1.616 (0.728)<br><br>1.287 (0.591) | 1.104 (0.994)<br><br>1.045 (0.884) |
| Minimal rejections (Delorme) | 1.519 (0.771)<br><br>1.234 (0.693) | 1.149 (1.061)<br><br>1.153 (1.057) |

**Supplementary Table S4.** Root mean square error values between the averaged ERP within each participant for each condition separately after applying different extreme artifact electrode and period rejections using either the default RELAX wICA settings (measured at FPz for the Go/Nogo dataset) or the targeted wICA settings which only addressed blinks and muscle activity (measured at FP2 for the N400 dataset).

One potential caveat to these results is that extreme rejection settings for the Go/No-go dataset were tested using the default wICA, which our primary results demonstrated could inflate effect sizes because of imperfect source separation. However, our test of the RMSE between the ERPs before and after artifact reduction with the wICA default approach at a fronto-polar electrode suggested that the different extreme rejection settings were not associated with differences in the distortion of the ERP in the Go/No-go dataset (see Supplementary materials Table S4, Section 1). Additionally, when we tested the targeted wICA cleaning approach in the N400 dataset, our results indicated that the RELAX moderate extreme artifact rejection approach was associated with the least distortion of ERPs (see Supplementary materials Table S4, Section 1), performing significantly better than the approach suggested by Delorme (2023). As demonstrated by our primary analyses reported in the main manuscript, targeted wICA restricts artifact cleaning to only eye-movement artifact periods and muscle activity frequencies, so the enhanced performance of the moderate extreme artifact rejection approach is unlikely to be due to the distortion of ERPs by component rejection. Instead, we suspect this result is caused by the less aggressive extreme rejection approaches including more high amplitude artifacts that exceed the thresholds for cleaning within the targeted wICA function, while not necessarily exceeding the criteria we set to exclude the raw epochs for the analysis of RMSE. As a result, these extreme artifacts may have influenced the raw ERP, and were cleaned by the targeted wICA, leading to a larger divergence between the raw and cleaned ERP and a higher RMSE.

In contrast, the moderate extreme rejection settings may have excluded the EEG periods and electrodes that would contribute to the threshold being exceeded for targeted wICA to apply cleaning, leading to an increased match between the raw ERP and cleaned ERP. The effect on our results of including these extreme artefacts is uncertain, as our metrics do not capture their impact. As such, we cannot determine the effect of removing these artifacts prior to ICA decomposition in contrast to including and cleaning these extreme artifacts. However, given that these artifacts are selected based on the application of statistics to detect "extreme" outliers, we suggest it is better to assume they do not reflect retrievable neural activity and to exclude them, rather than assume they can be effectively cleaned without adversely affecting the performing of the artifact reduction methods. This approach of excluding more initial extreme artifacts does exclude more of the data, but it also reduces the differences between the pre-cleaned and post-cleaned data without leaving more artifacts in the data, an outcome that might be considered preferable and in alignment with a philosophy that minimally manipulating the data prior to analysis is desirable.

Overall, given that a moderate to light stringency of extreme artifact rejection settings was associated with a larger between condition effect size, and the moderate stringency approach was associated with the least distortion of the ERP in the N400 dataset, we applied moderate stringency of extreme artifact rejection in our primary analyses. We also tentatively recommend that future research apply a moderate to light stringency of extreme artifact rejection settings (Bailey, Biabani, et al., 2023; Bigdely-Shamlo, Mullen, Kothe, Su, & Robbins, 2015; Nolan, Whelan, & Reilly, 2010). However, further research dedicated specifically to this issue is needed to explore extreme artifact rejection settings prior to ICA in more detail to determine the optimal approach.

***Section 2: Comparison pipelines***

To determine the effects of component-based artifact reduction methods, we tested a range of pipelines after applying band-pass filtering the data using a fourth order Butterworth filter from 0.5 to 80Hz, and notch filtering from 47 to 53Hz for the Go Nogo dataset and 57 to 63Hz for the N400 dataset. We then applied the RELAX moderate stringency electrode and extreme artifact period rejections described in the previous section. Following the filtering and extreme artifact rejection steps, our targeted wICA method and eight comparator pipelines were tested. These included: 1) ICA subtract, 2) ICA subtract light, 3) wICA, 4) IVA, 5) MWF, 6) another novel method that we tested: canonical correlation analysis to clean muscle activity followed by generalised eigenvector decomposition to clean blinks (CCA GED), 7) DSS, and 8) a regression blink reduction method. The ICA subtract, ICA subtract light, and wICA methods all used the PICARD algorithm to decompose data (Frank, Makeig, & Delorme, 2022). The ICA subtract, ICA subtract light, wICA, and IVA methods all selected artifact components for reduction using ICLabel. Artifact components were identified when the classification likelihood was maximum for an artifact category by ICLabel for the ICA subtract, wICA, and IVA methods. A 0.8 likelihood threshold of being a blink or muscle component was used for the ICA subtract light pipeline. Artifact components were replaced with zeros prior to reconstruction into the scalp space for the ICA subtract, ICA subtract light, and IVA pipelines. For the wICA pipeline, a wavelet transform was applied to the artifact components in an attempt to characterise the artifact contribution to the component time-series, which was then subtracted from the original component, before the data were reconstructed back into the source space (Bailey, Hill, et al., 2023).

The independent vector analysis (IVA) pipeline involved applying the IVA algorithm developed by Anderson, Adali, and Li (2011) to a matrix of the EEG data with a single delay embedding version of that same EEG data, with the delay embedding separated from the original data by 8ms. This approach has been reported to lead to excellent separation of blink and muscle artifacts (Barban,

Chiappalone, Bonassi, Mantini, & Semprini, 2021). In particular, the inclusion of the autocorrelation in the IVA algorithm makes it highly effective at separating muscle components (Barban et al., 2021).

MWF was implemented by obtaining a template of the artifact and non-artifact periods in the data, then applying the MWF algorithm to characterize and clean the artifact periods (Somers, Francart, & Bertrand, 2018). This was performed with artifact periods being identified using the RELAX default settings (Bailey, Biabani, et al., 2023). A delay period of 8 samples was implemented, meaning that for the muscle artifacts, the delay embedded matrix used for MWF cleaning spanned 17 samples (with delay embeddings constructed every sample for 8 samples on either side of the original data). However, when cleaning blink artifacts, we implemented a delay spread, whereby each delay embedding was separated from its neighbouring delay embedding by 16ms (rather than 1ms, as would be the case with MWF's traditional application when data are sampled at a commonly used 1000Hz sampling rate). This characterised a full 272 samples (and ~272ms given our 1000Hz sampling rate for our Go Nogo dataset and 1024Hz sampling rate for the N400 dataset). Our informal testing indicated that characterising this broader period provided superior blink artifact cleaning compared to the original version of the MWF with delay embeddings every consecutive sample. Our primary results indicated that MWF with a sparsely separated delay embedding showed the (distant) second best performance at protecting the ERP waveform from distortion by cleaning (with targeted wICA performing the best). As such, if researchers are concerned about the number of algorithm degrees of freedom involved in our targeted wICA cleaning method, using a delay spread version of the MWF approach might be the next best option. To enable researchers to use this method, we have made it possible to set staggered delay embeddings within the MWF cleaning approach in the RELAX pipeline. Users should note that the MWF application within RELAX relies on the MWF algorithm introduced by Somers, Francart and Bertrand (2018), which should be cited if the MWF cleaning approach within RELAX is used.

We also tested the denoising source separation (DSS) method. To achieve this, we first separated the data into 4 second epochs time-locked to the Go and Nogo stimuli, then applied polynomial robust detrending with an order of 2 rather than band-pass filtering (as robust detrending is applied prior to DSS by the author of the method - de Cheveigne (2023)). We also cleaned the line noise prior to DSS using Zapline (de Cheveigné, 2020). Then, in an initial approach to address blink activity, periods within high amplitude components detected by DSS that exceeded 3SD were replaced with the 3SD value, as recommended within the DSS approach. Next, data were cut into shorter epochs (-100 to 1000ms surrounding stimuli) and cleaned with a repeat-biased DSS, keeping 10 components (de Cheveigné & Parra, 2014). This repeat-biased DSS separates the data into components using the repeated ERP information within the trials, principal component analysis, and a bias filter to obtain components that involve enhancements of the power from the neural signal of interest and reductions of power from the noise sources (which includes artifacts). These components are ranked by the ratio of the signal power to the noise power, and components below a specific signal to noise ratio can be deleted (components above 10 in the case of our data). This approach has been suggested to enhance signal to noise ratios while relying less on potentially inconsistent modelling (as is the case for the blind source separation methods such as ICA, wICA, and IVA) (de Cheveigné & Parra, 2014).

Next, we tested an additional component subtraction method that we developed in an initial attempt to address the imperfect source separation issue, which we refer to as CCA GED. This pipeline first applied an extended canonical correlation analysis to address muscle activity (Janani et al., 2018) then applied a generalised eigenvector decomposition (GED) to address blink activity. To achieve this, the pipeline used blink periods (identified by RELAX) to construct a signal covariance matrix and non-blink periods to construct a reference covariance matrix (Cohen, 2022). We then performed a GED to decompose data into components reflective of the maximal difference between the blink and non-blink periods. Component time-series that contained blink artifacts were identified as showing absolute amplitudes during blink periods that were significantly larger than non-blink periods, tested using a one-way t-test of the ratio between the absolute amplitude within each blink period and the absolute amplitude of the overall data excluding blink periods, with the t-test testing for differences between this ratio and a value of 1. Components identified as reflecting blink artifacts were then replaced with zeros before scalp space data were reconstructed.

Finally, for the N400 dataset that provided a VEOG electrode under the right eye, we tested an artifact aligned regression-based blink cleaning method, which involves subtracting beta weights from each electrode rather than subtracting an artifact component. To achieve this, we constructed a virtual VEOG electrode by subtracting the signal from the electrode under the right eye from the signal at FP2. We then applied the artifact aligned average method, averaging the data aligned to each blink maximum (detected by RELAX) to obtain a representation of the average blink activity at the reconstructed VEOG electrode and each EEG electrode. This artifact aligned average was used to obtain the beta weights with a least squares regression for the effect of the blink (measured with the virtual VEOG channel) on each EEG electrode, which were then used to correct the blink artifact in the continuous data (Croft & Barry, 2000).

**Section 3: Supplementary Results – Muscle Artifact and Other Artifact Components**

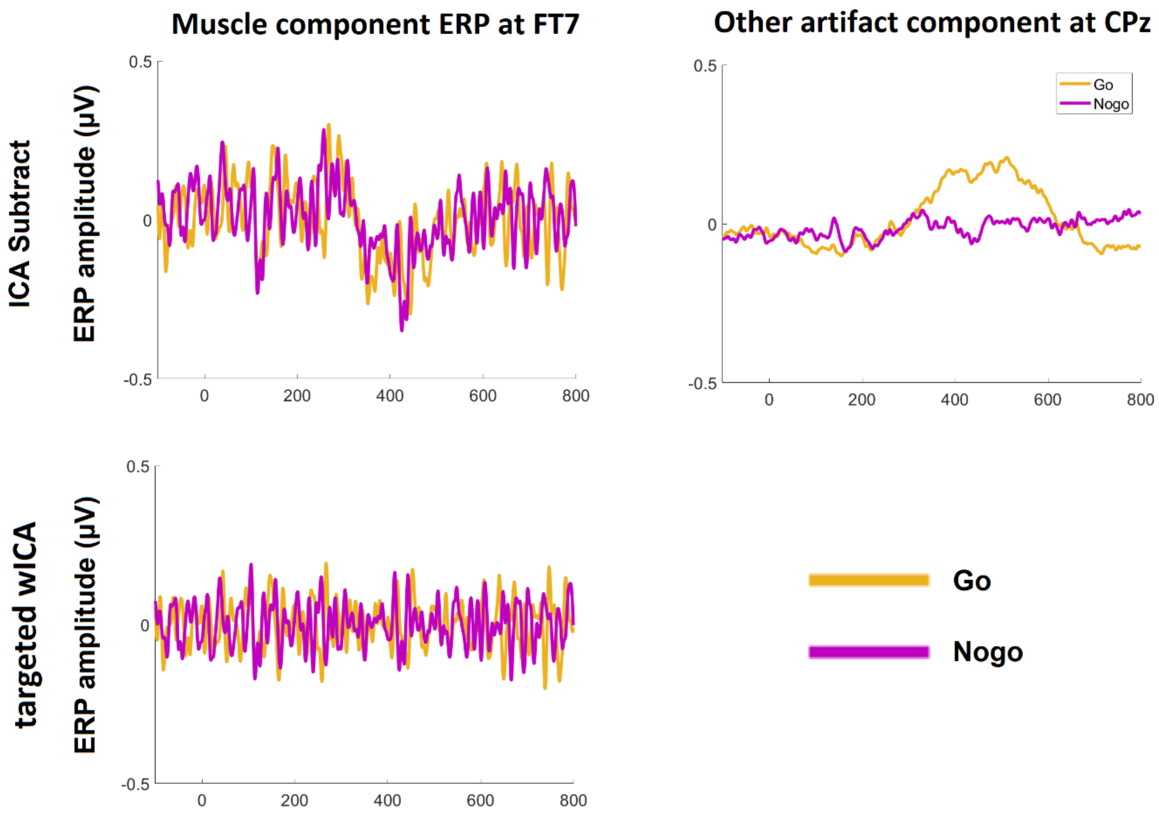


Figure S1. Left: Event-related potentials (ERP) from the Go Nogo dataset at the FT7 electrode obtained from components that were identified as muscle artifacts using the ICA subtract method being projected back to the scalp space data only (top), and the artifact signal that was removed by the targeted wICA method (bottom). Note that because the targeted wICA method high-pass filtered the muscle artifact components at 15Hz before subtracting the muscle artifact components from the data, contribution from the P3 ERP is no longer mixed into the muscle artifact estimate, and the P3 ERP contained in the muscle artifact component is therefore preserved in the data. In contrast, the ICA subtraction method removed the P3 ERP contribution along with the muscle activity. Although in this specific dataset, the P3 contribution captured within the muscle component was small in amplitude and did not differ between conditions (so its subtraction would have no effect on between condition comparisons at the scalp space), it is possible that the ERP caught within the muscle components might differ in other datasets, affecting between condition comparisons. Right:

The Go Nogo ERP constructed from the projection to scalp space data from only components that were identified as artifacts other than muscle or eye movements. Note that an ERP signal is mixed into these artifact components and is visible from 400 to 600ms post stimuli. Although this ERP signal mixed into the artifact component is only small in amplitude, the signal differed between the Go and Nogo conditions. As such, an artifact subtraction method that removed these artifact components would affect the scalp space comparisons. Given this result, and the difficulty with characterising and specifically reducing non-eye-movement and non-muscle artifacts, our targeted wICA approach does not reduce these additional artifact types.
